# Initial Learning in the Brain: From Rules to Action

**DOI:** 10.1101/2024.03.12.584566

**Authors:** Sofia Fregni, Uta Wolfensteller, Hannes Ruge

**Author notes:** Correspondence concerning this article should be addressed to Sofia Fregni, Bürogebäude Zellescher Weg 17, 01069 Dresden. The authors made the following contributions. Sofia Fregni: Conceptualization, Formal Analysis and Investigation, Writing - Original Draft Preparation, Writing - Review & Editing; Uta Wolfensteller: Conceptualization, Writing - Review & Editing; Hannes Ruge: Conceptualization, Writing - Review & Editing, Supervision.

## Abstract

We used fMRI to investigate the neural changes and representational dynamics associated with different learning modes during initial learning and subsequent implementation of previously acquired stimulus-response (S-R) associations. We compared instruction-based learning (INS) and trial-and-error learning (TE) via a third observation-based learning (OBS) condition. This was yoked to the TE condition and shared features with both, the INS and TE conditions. During learning, neural changes were observed in the Frontoparietal and Default Mode Networks across learning modes, consistent with a general decrease in cognitive control demand as learning progresses. INS and TE exhibited condition-specific signal changes, which we interpreted in the context of covert motor preparation during INS, and intentional action and increased cognitive control demand during early TE trials, respectively. Multivariate pattern analysis revealed individual rule information in bilateral prefrontal, premotor, and parietal cortices across learning modes. Most regions revealed consistent representations of individual S-R rules between the learning stage and subsequent implementation stage, regardless of the learning mode. This suggests that initially formed S-R rule representations guide task performance during S-R rule implementation, irrespective of how they are acquired. Finally, within the primary motor cortex, individual S-R rules were decodable during the learning stage not only when motor responses were overtly executed, as in TE, but also in the absence of overt motor execution, as in INS. This finding substantiates previous claims of covert motor preparatory mechanisms during INS.

---

1 **Introduction**

Goal-directed behavior is a fundamental aspect of human experience, as it enables us to flexibly navigate the environment, and design complex plans to achieve our goals. Laboratory settings offer an avenue to study it by creating simple goals and motivating participants to attain them through more or less direct rewards, such as monetary outcomes or performance feedback. Achieving our desired outcomes in a novel environment via trial-and-error learning, which involves repeated attempts, contingent feedback, and necessary adjustments, is, however inherently slow and error-prone. Instruction-based learning, on the other hand, refers to the unique human capacity to rapidly and efficiently learn via explicit instructions. In laboratory settings, it has been shown that humans can successfully learn newly instructed S-R links instantaneously and achieve a high degree of fluency in the space of a few practice trials (Monsell & Graham, 2021; Ruge & Wolfensteller, 2010; Wolfensteller & Ruge, 2012). It follows that, at least at the behavioral level, instruction-based learning bears an indisputable advantage compared to trial-and-error learning strategies (Li et al., 2011; Monsell & Graham, 2021; Ruge, Karcz, et al., 2018).

One potential explanation for the learning advantage of explicit instructions, compared to trial-and-error, might hinge on distinct working memory (WM) processes at play during either learning mode. Ruge, Karcz, et al. (2018) directly compared instruction-based to trial-and-error learning at the behavioral level. Interestingly, a positive correlation between WM-span scores, which measure *declarative* WM (cf. Lee et al., 2020), and accuracy was found in trial-and-error learning, especially in early trials, but not in instruction-based learning. From a theoretical standpoint, it is plausible that trial-and-error learning heavily relies on declarative WM (Oberauer, 2009, 2010) during rule exploration, at least as long as the process of rule extraction is not completed (Monsell & Graham, 2021; Ruge, Karcz, et al., 2018). Accordingly, the agent learning via trial-and-error needs to recognize and recall previously encountered S-R links and response outcomes to extract the correct S-R association (Collins & Frank, 2013; Mohr et al., 2018). For actual S-R rule implementation, however, be it correct or incorrect, it is plausible that each S-R rule is represented procedurally (Monsell & Graham, 2021; Ruge, Karcz, et al., 2018). Yet, different from learning via instruction, during trial-and-error learning, both declarative and procedural S-R rule representations would need to be updated after each negative feedback until the correct mapping has been implemented. Evidence comes from a recent study by Monsell and Graham (2021) where participants engaged in a rapid learning task involving novel one-to-one S-R links that could be learned via instruction or trial-and-error. The hypothesis was that novel instructed tasks might be initially represented declaratively, i.e. what the instruction requires, and later transformed into executable programs of the like “If X, press right; if Y, press left”. To explore the shift from declarative to procedural representation of the S-R links, the authors used between-stimulus phonological similarity as an index of declarative WM. Interestingly, phonological similarity negatively affected performance during the implementation of previously instructed S-R mappings only for a few practice trials, whereas it took three times the amount of practice trials to reach the same asymptotic performance level in the trial-and-error manipulation. This favors the hypothesis that trial-and-error taps into declarative WM more than instruction-based learning and that the latter encourages a faster proceduralization during the implementation of previously instructed rules (cf. Formica et al., 2020; González-García et al., 2021). Complementary insights come from studies showing that merely instructed (i.e., never overtly implemented) S-R rules can negatively impact behavioral performance when these are incongruent with the response requirements in a subsequent task (e.g. Braem et al., 2019; Brass et al., 2009; Entel et al., 2014; Liefooghe et al., 2012; Liefooghe & De Houwer, 2018; Wenke et al., 2015). This, in turn, suggests that rule proceduralization during instruction-based learning might take place as early as before rule implementation, i.e. in the absence of overt response execution (González-García et al., 2021; see also Meiran et al., 2015; Palenciano et al., 2021). Brass et al. (2017) proposed that this early translation of novel instructions into condition-action rules might involve mental imagery and covert S-R rule implementation during the instruction phase to guide subsequent overt response execution in the implementation stage (González-García et al., 2021; cf. Palenciano et al., 2021). This is supported by neuroimaging studies that found motor and premotor activity during the instruction period, which is detached from any overt motor response implementation (Hampshire et al., 2019; Hartstra et al., 2011; Ruge & Wolfensteller, 2010). Additionally, Muhle-Karbe et al. (2017) compared how the same S-R instructions were represented in the brain when these had to be implemented or recalled later on. Instructions were found to be decodable in an extended bilateral frontoparietal network during the delay period following instruction only in implementation blocks (cf. González-García et al., 2021). Furthermore, decoding accuracy in visual regions during the delay positively correlated with RTs during instruction implementation, suggesting that preparatory mechanisms took place in associating the visual stimuli to their response when these had to be implemented later on. Last, the authors compared the similarity between instruction-related patterns during the instruction and the delay period in frontoparietal regions and found greater similarity in memorization vs. implementation blocks, as well as between the instruction period of implementation blocks and the delay period of memorization blocks. This suggests that simple to-be-implemented S-R instructions are rapidly represented in frontoparietal regions as early as during the instruction period and that the same type of information from when it is instructed to the period following instruction but preceding implementation goes through some kind of transformation.

The central goal of the present study was to investigate the neural bases underlying the initial phase of learning by instruction and trial-and-error via the use of fMRI. In particular, we hypothesized differential involvement of declarative and procedural WM, with a faster proceduralization of rules that were learned via instruction as opposed to rules that were learned via exploration. Studying the initial phase of learning was of particular interest as this contrasted with previous studies examining time or trial bins of implementation trials following instruction to characterize the proceeding of learning (Hampshire et al., 2016, 2019; Ruge et al., 2019). In our study, we distinguished initial learning from subsequent practice, thereby preventing the possible conflation of the two phases.

In summary, the hypothesis that merely instructed S-R rules are rapidly stored in procedural WM before their implementation could explain the very fast rate at which this type of learning leads to task automatization (Baumann et al., 2023; González-García et al., 2021; Mohr et al., 2016; Ruge & Wolfensteller, 2010; Wolfensteller & Ruge, 2012) and the performance advantage compared to trial-and-error strategies (Monsell & Graham, 2021; Ruge, Karcz, et al., 2018). However, while behavioral evidence suggests that in trial-and-error and instruction-based learning the extent of declarative and procedural WM engagement might differ, neuroimaging studies point towards common neural dynamics. To elaborate, common activation clusters supporting human value-based learning and decision-making in the brain encompass regions that have been found to increase in activity from the early implementation of previously instructed S-R rules (Ruge & Wolfensteller, 2010, 2013), as well as during the delay following first-time instruction (Muhle-Karbe et al., 2017). These regions include frontal and parietal cortices, as well as the ventral striatum, pre-SMA, and anterior insula (Liu et al., 2011). In addition, there is evidence for similar neural dynamics underlying trial-and-error and instruction-based learning. These include a frontoparietal decrease and Default Mode Network (DMN) activity increase (e.g. Bédard & Sanes, 2009; Hampshire et al., 2016, 2019; Mohr et al., 2016; Ruge & Wolfensteller, 2010, 2013), as well as an activity increase in the ventral striatum (Ruge et al., 2019; Ruge & Wolfensteller, 2010, 2015), known for its sensitivity to various types of rewards, including positive feedback (for an overview, Daniel & Pollmann, 2014). Yet, the lack of a direct comparison between the two learning modes leaves a window open to the possible differences and commonalities. It follows that it would be extremely informative to investigate how learning unfolds trial-by-trial, at each S-R rule encounter, under instruction and trial-and-error. A study by Ruge et al. (2019) combined the approach to track learning-related neural changes on a trial-by-trial basis with multivariate analysis to identify brain activity patterns specific to individual S-R rule identities (e.g., ‘house-right index finger’; ‘flower-right middle finger’) during the initial implementation stage after first-time instruction. Identity-specific S-R rule representations were found in the left ventrolateral prefrontal cortex (VLPFC) as early as from the first implementation trial after instruction. The focus on implementation trials, however, raises questions about whether and how these brain representations are already built during preceding instruction trials. With this in mind, we employed a task design similar to Ruge et al. (2019), but novel in the way it allowed us to broaden the focus to the study of the initial phase of learning, prior to actual S-R rule implementation. Via univariate and multivariate analysis of fMRI data, we tracked the neural changes and representational dynamics of S-R rules while being instructed vs. explored via trial-and-error. To be able to compare the fundamentally different instruction-based and trial-and-error learning modes, we included a third learning condition of interest, termed observation-based learning. In this condition, learning required the mere observation of S-R rules, like in instruction-based learning. However, rules could be correct or incorrect, as indicated by post-rule feedback, requiring rule extraction, akin to trial-and-error-learning. Finding an effect specific to trial-and-error but not instruction- and observation-based learning would link the effect to motor implementation, whereas an effect common to trial-and-error and observation but not to instruction would link the effect to error processing. In addition, we aimed to examine how individual S-R rules might be differentially represented in the learning and implementation stages and whether these representations stay consistent from one stage to the other. This latter investigation covered several regions of interest (ROIs) that had previously been identified as underlying instruction-based and trial-and-error learning in the existing literature. These included the ventral and dorsal LPFC (Ruge et al., 2019), the inferior and superior parietal cortices (e.g. Bode & Haynes, 2009; Cole et al., 2010; Ruge & Wolfensteller, 2010, 2013), as well as left motor and bilateral premotor cortices (Hampshire et al., 2019; Hartstra et al., 2011; Ruge & Wolfensteller, 2010). The inclusion of the premotor and motor cortex in the set of ROIs is of particular relevance to our hypothesis that mere instruction trials would already involve covert motor preparation, possibly mirroring an early proceduralization of rules while being instructed. Last, this complements recent findings by Ariani et al. (2022), demonstrating the utility of multivariate compared to univariate analysis in dissociating neural effects of covert motor planning from overt motor execution.

## 2 Methods

The study was pre-registered on AsPredicted (https://aspredicted.org/9q9m6.pdf). Details and changes with respect to the pre-registration are in the [Supplementary material].

Group-level images, as well as ROI-based MVPA and behavioral data are available at https://osf.io/5mcf8/.

### 2.1 Participants

A total of 84 German native speakers participated in the experiment. Data from 4 participants were excluded due to technical errors during the experiment. The final data set included 80 participants (53 females and 27 males, mean age = 23.92, SD = 4.37, range 18-34). Participants did not have a previous history of neurological or psychiatric disorders, had normal or corrected-to-normal vision, and were right-handed. All provided written informed consent prior to the experiment start and received payment or university credits for their participation in the study. The experimental protocol was previously approved by the Ethics Committee of the Technische Universität Dresden (EK586122019).

### 2.2 Experimental material and task

The task was a rapid learning task (Figure 1a) that required participants to learn novel S-R links in the space of a few trials. In order to track learning processes in a trial-by-trial manner, each experimental block involved a set of 4 different novel stimuli that were coded in terms of stimulus repetitions 1 to 8. Within each block, each stimulus was repeated 4 times during the learning stage (stimulus repetitions 1 to 4), and 4 times during the implementation stage (stimulus repetitions 5 to 8). And it was linked to a unique response that participants had to learn via instruction, trial-and-error, or observation of the S-R links. Each experimental block started with a general instruction cue that informed participants about the upcoming learning condition. This was either to remember, try, or observe the S-R links (in German *Merken, Ausprobieren* or *Beobachten*) in the instruction-based, trial-and-error, and observation-based learning conditions, respectively. During the learning stage, each stimulus was presented at the center of the screen above the schematic drawing of 4 buttons. These were represented by 4 rectangles, one next to the other, one for each possible response. In the instruction-based learning condition, each stimulus was correctly paired to its response, i.e. button press. The correct response that was associated with each stimulus was cued by the correspondent button on the screen that differed from the others for its black vs. white color (Figure 1a, Learning stage, instruction). In the trial-and-error learning condition, participants were asked to respond to stimuli via a button press with the index, middle, ring, or pinky finger of the right hand. At the time of the button press, the correspondent button on the screen turned black, providing participants with visual feedback about their response (Figure 1a, Learning stage, trial-and-error). After each trial, correct or incorrect feedback was presented depending on whether the participant pressed the correct or incorrect button, respectively. In the observation-based learning condition, participants were passively attending the presentation of S-R links that could be correct or incorrect, as indicated by the following feedback (Figure 1a, Learning stage, observation). In this sense, this learning condition resembled the trial-and-error learning condition in the measure of incorrect feedback processing. However, just like in the instruction-based learning condition, participants were not required to respond to trials during the learning stage, but only to observe what was presented on the screen. Each stimulus was presented together with the schematic drawing of 4 white buttons underneath. The identity of the observed response was cued by one of the 4 buttons on the screen that turned black after each stimulus presentation. Based on whether the observed S-R link was correct or incorrect, each trial was followed by correct or incorrect feedback, respectively.

**Figure 1.**
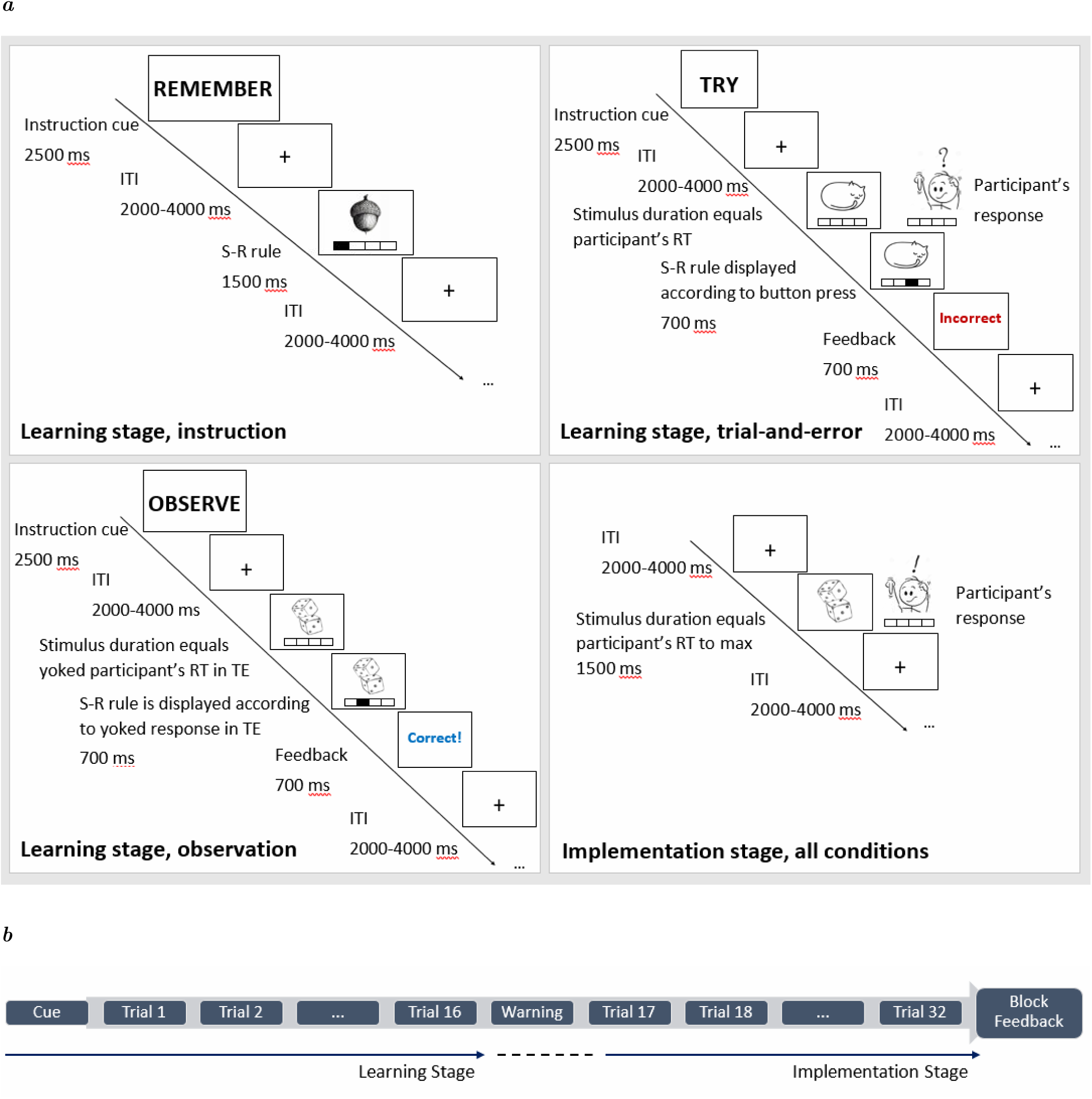
a) The figure depicts an exemplary learning trial for each learning condition, as well as an examplary implementation trial (same across conditions). ITI = Inter-trial interval. b) Trial structure per each of the 24 experimental blocks. Each block comprised a total of 32 trials, 16 per each stage. Each block started with a cue, referring to the learning condition cue. After the first 16 learning trials (learning stage), a warning message reminded participants of the start of the implementation stage for the next 16 trials. At the end of each block, the corresponding block accuracy feedback was provided.

Response onsets and response identity in the observation-based learning condition were built upon the trial-and-error learning condition as follows. Data were collected in sets of 5 participants, where the response to each trial in the trial-and-error condition was analyzed in terms of RT and accuracy (i.e. the number of correct and incorrect trials) per each individual subject. The next 5-participant set in the observation-based learning condition included trial-by-trial feedback after the observation of each S-R link that resembled the feedback that each individual participant received in the trial-and-error condition in the previous 5-participant set. This created a closed, well-controlled loop between the observation-based and trial-and-error learning conditions, as well as more naturalistic response patterns in the observation-based learning condition. Response identities and onsets in the observation-based learning condition for the first 5 participants were constructed based on previous behavioral pilot data.

The implementation stage was structured in the same way for all learning conditions and consisted of the same stimuli as in the correspondent learning stage (Figure 1a). Participants were required to respond to trials via a button press with the index, middle, ring, or pinky finger of the right hand. No trial-by-trial feedback was presented at this stage of the experimental procedure.

The stimuli consisted of 108 black and white drawings representing various animated and inanimate objects. These were distributed across 24 experimental blocks, each consisting of 4 stimuli, plus 3 additional blocks designated for practice before the start of the experiment. The assignment of stimuli to blocks and the order of stimuli and block presentation were pseudo-randomized across participants. Importantly, stimuli sequences were built in the form of stimulus repetition chunks, where participants attended all the stimuli for the first time (stimulus repetition 1), then for the second time (stimulus repetition 2), etc. . . until stimulus repetition 8. Following up on earlier work (Ruge, Legler, et al., 2018), in the present study we used a refined trial sequencing scheme to eliminate bias in the planned MVPA analysis for any combination of stimulus repetitions. Different from the original version (Ruge, Karcz, et al., 2018; Ruge et al., 2019), this modified approach enabled us to compute pattern similarities during both the learning and implementation stages. This allowed us to test the consistency of rule representations across the different stages of the experiment.

### 2.3 Experimental procedure

The experiment took place at the Neuroimaging Center in Dresden. Before the actual experiment, participants were familiarized with the task outside the scanner. The procedure for the practice and the real experiment were the same. The practice consisted of one block of 32 trials (4 stimuli x 8 repetitions) per each experimental condition. The 12 stimuli, 4 per condition, that were used during the practice were not re-used in the real experiment. Before the practice, participants attended a few slides with precise instructions about the task and the S-R links presentation. At this stage, participants learned to associate each finger press/response to the correspondent button that was drawn onto the screen underneath each stimulus. Following the practice, participants moved into the scanner and started the real experiment. The full experiment consisted of 24 blocks with 32 trials per block (Figure 1b). These were administered in the form of 4 functional runs, each lasting 20 minutes and consisting of 6 blocks, with 2 blocks per condition. Block onsets within the same run were jittered with an interval of 2 or 4 seconds. Blocks were randomized across participants such that the same run for different participants always consisted of different blocks. Participants were provided with ear plugs to prevent ear damage due to the scanner noise for the whole duration of the experiment. Experimental material was projected onto the screen that is placed in the back of the scanner so that participants could see through the rear-facing mirror. Each experimental block started with a fixation cross that was presented at the center of the screen for 2 seconds. This was followed by a 2.5-s instruction cue that instructed participants on the upcoming learning condition. Each trial lasted a maximum of 1500 ms in all conditions or until the onset of a response in the observation-based and trial-and-error conditions. In observed trials, when a response was observed, and in trial-and-error trials, when a response was made, the button on the screen that corresponded to the button pressed in the scanner turned black for 700 ms to give visual feedback to the participants (Figure 1a, Learning stage, observation, and Learning stage, trial-and-error, respectively). This was followed by a *Korrekt!* or *Falsch* feedback depending on whether the response was correct or incorrect, respectively. Accuracy-based feedback during learning in the trial-and-error and observation-based learning conditions lasted 700 ms and was depicted in blue for correct trials and red for incorrect trials. When no response was detected in the trial-and-error condition, a 700-ms red warning message appeared on the screen to inform participants. If participants pressed a button to respond to trials during the learning stage in the instruction- and/or observation-based learning conditions, where a response was not required, a message appeared at the center of the screen for 2.5 seconds asking them not to respond to trials and reminding them about the present learning condition. Trials were interleaved by a fixation cross that was presented at the center of the screen and jittered between 2 and 4 seconds. After 16 trials, a 2-s message was presented at the center of the screen, warning participants about the upcoming implementation stage. This consisted of another set of 16 trials, where no trial-by-trial feedback was provided. The absence of a response in implementation trials triggered a 500-ms red message warning participants that no response was detected. Like in the learning stage, the inter-trial interval was jittered between 2 and 4 seconds and trials stayed on screen until a response was made or for a maximum of 1500 ms. Each experimental block ended with individual performance-based feedback where block accuracy in the implementation stage was depicted as a percentage of correct trials in that individual block. Block-specific feedback was presented on screen for 3 seconds. At the end of the experiment, the total accuracy for the whole experiment was presented on the screen for 4 seconds. Participants were previously informed that with a total accuracy above 85% of correct trials in the whole experiment, they would have received extra money. The performance-based extra payment entailed 1 extra euro for accuracy between 85 to 89%, 2 extra euros for 90 to 94% accuracy, and 3 extra euros for overall performance above 95% accuracy. The task was coded and presented via E-Prime 3.0 software (Psychology Software Tools, Pittsburgh, PA; RRID:SCR_009567). Stimuli sequences and experimental lists were created in Matlab (version R2020b, The Math Works, Inc., 2020) and RStudio (R Core Team, 2021).

### 2.4 Data acquisition

MRI data were acquired in a Siemens MAGNETOM 3T Trio Tim scanner with a 32-channel head coil. The BOLD sequence we used allowed for a good frontal resolution and minor ghosting effect (TR = 2070 ms, TE = 25 ms, flip angle = 80°, FoV = 192 mm, voxel size 3.0×3.0×3.2 mm, EPI factor = 64). Slices were acquired in descending sequential order for a total of 36 slices per volume. Each functional acquisition lasted about 25 minutes and it was followed by a GRE EPI fieldmap sequence with the same voxel size as the bold sequence in order to correct for field inhomogeneities of the bold acquisition (TR = 439 ms, TE 1 = 5.32 ms, TE 2 = 7.78 ms, flip angle = 45°, FoV = 192 mm, voxel size 3.0×3.0×3.2 mm). At the end of the 4 functional runs, a high-resolution T1-weighted anatomical image with full brain coverage was acquired for each participant (MPRAGE, TR = 1900 ms, TE = 2.26 ms, flip angle = 9°, FoV = 256 mm, voxel size = 1.0×1.0×1.0 mm).

### 2.5 Data processing

Raw MRI data were converted into BIDS 1.2.1 (Gorgolewski et al., 2016, RRID:SCR_016124) format before preprocessing.

Preprocessing was performed using *fMRIPrep* 21.0.0 (Esteban, Markiewicz, et al., 2018; Esteban, Blair, et al., 2018, RRID:SCR_016216), which is based on *Nipype* 1.6.1 (Gorgolewski et al., 2011; Gorgolewski et al., 2018, RRID:SCR_002502).

#### 2.5.1 Preprocessing of B0 inhomogeneity mappings

A total of 4 fieldmaps per subject were used within the input BIDS structure. A *B0* nonuniformity map (or *fieldmap*) was estimated from the phase-drift map(s) measure with two consecutive GRE (gradient-recalled echo) acquisitions. The corresponding phase-map(s) were phase-unwrapped with prelude (FSL 6.0.5.1:57b01774).

#### 2.5.2 Anatomical data preprocessing

Each participant’s T1-weighted (T1w) image was corrected for intensity non-uniformity (INU) with N4BiasFieldCorrection (Tustison et al., 2010), distributed with ANTs 2.3.3 (Avants et al., 2008, RRID:SCR_004757), and used as T1w-reference throughout the workflow. The T1w-reference was then skull-stripped with a *Nipype* implementation of the antsBrainExtraction.sh workflow (from ANTs), using OASIS30ANTs as target template. Brain tissue segmentation of cerebrospinal fluid (CSF), white-matter (WM) and gray-matter (GM) was performed on the brain-extracted T1w using fast (FSL 6.0.5.1:57b01774, RRID:SCR_002823, Zhang et al., 2001). Volume-based spatial normalization to one standard space (MNI152NLin2009cAsym) was performed through nonlinear registration with antsRegistration (ANTs 2.3.3), using brain-extracted versions of both T1w reference and the T1w template. The following template was selected for spatial normalization: ICBM 152 *Nonlinear Asymmetrical template version 2009c* (Fonov et al., 2009, RRID:SCR_008796, TemplateFlow ID: MNI152NLin2009cAsym).

#### 2.5.3 Functional data preprocessing

For each of the 4 BOLD runs per subject (across all tasks and sessions), the following preprocessing was performed. First, a reference volume and its skull-stripped version were generated using a custom methodology of *fMRIPrep*. Head-motion parameters with respect to the BOLD reference (transformation matrices, and six corresponding rotation and translation parameters) are estimated before any spatiotemporal filtering using mcflirt (FSL 6.0.5.1:57b01774, Jenkinson et al., 2002). The estimated *fieldmap* was then aligned with rigid-registration to the target EPI (echo-planar imaging) reference run. The field coefficients were mapped on to the reference EPI using the transform. BOLD runs were slice-time corrected to 1.01s (0.5 of slice acquisition range 0s-2.02s) using 3dTshift from AFNI (Cox & Hyde, 1997, RRID:SCR_005927). The BOLD reference was then co-registered to the T1w reference using mri_coreg (FreeSurfer) followed by flirt (FSL 6.0.5.1:57b01774, Jenkinson & Smith, 2001) with the boundary-based registration (Greve & Fischl, 2009) cost-function. Co-registration was configured with six degrees of freedom. The BOLD time-series were resampled into standard space, generating a *preprocessed BOLD run in MNI152NLin2009cAsym space*. First, a reference volume and its skull-stripped version were generated using a custom methodology of *fMRIPrep*. All resamplings can be performed with a *single interpolation step* by composing all the pertinent transformations (i.e. head-motion transform matrices, susceptibility distortion correction, and co-registrations to anatomical and output spaces). Gridded (volumetric) resamplings were performed using antsApplyTransforms (ANTs), configured with Lanczos interpolation to minimize the smoothing effects of other kernels (Lanczos, 1964). Non-gridded (surface) resamplings were performed using mri_vol2surf (FreeSurfer).

Many internal operations of *fMRIPrep* use *Nilearn* 0.8.1 (Abraham et al., 2014, RRID:SCR_001362), mostly within the functional processing workflow. For more details of the pipeline, see the section corresponding to workflows in *fMRIPrep*’s documentation. ^1^

### 2.6 Data analysis

#### 2.6.1 Behavior

Behavioral analysis was implemented in R (R Core Team, 2021). Accuracy and response times (RTs) were analyzed in each condition in the implementation phase and in the trial-and-error learning stage. Data were subjected to repeated-measures ANOVAs (R package “afex,” Singmann et al., 2023) where accuracy or RT was the dependent variable as predicted by stimulus repetition for trial-and-error learning trials, and both stimulus repetition and learning condition for the analysis of implementation trials. Each model included the random factor subject. Results with p < .05 were further investigated via post-hoc estimated marginal means tests and pairwise comparisons (R package “emmeans,” Lenth, 2023) between stimulus repetition and condition depending on the model and the significant effects. Statistics of pairwise comparisons were adjusted for the number of tests via the multivariate t-distribution approach that is available within the R package “emmeans”.

#### 2.6.2 Univariate analysis

Preprocessed functional images were processed in SPM12 software package (Statistical Parametric Mapping; RRID:SCR_007037) and its implementation in Matlab (version R2020b, The Math Works, Inc., 2020). The first 3 scans for each run were discarded before starting image processing. Single-sub statistical maps were produced in the context of the General Linear Model, GLM. Correct and incorrect trials were modeled separately, excluding the very first trial in the learning phase and in the implementation phase. For both correct and incorrect trials, the original stimulus repetitions were re-coded based on accuracy (i.e., first correct/incorrect repetition, second correct/incorrect repetition, etc. . . ). Incorrect trials were modeled for the trial-and-error and observation-based learning conditions. For the learning phase of the instruction-based learning condition, all 16 trials were considered correct. The final GLM included regressors for the first trials of both the learning and implementation stages, instruction cues, implementation stage warning start message, as well as block feedback and sustained activity. Event-related model regressors were constructed by modeling each event as a stick function convolved with the canonical Hemodynamic Response Function, HRF. The model included a high-pass filter with a 200s cut-off.

For each subject, we created contrasts for the linear increase and decrease across stimulus repetitions in both the learning and the implementation stages, separately per each learning condition. In the context of implementation trials, we additionally implemented single-subject contrasts for the mean across repetition levels, separately for each learning condition. This was meant to test for the main effect of condition at the neural level, which was behaviorally observed uniquely in implementation trials. Before group-level statistics, single-subject contrast T-maps were smoothed with a 6 mm kernel and the SPM intra-cerebral volume mask was used to correct for voxels that lied outside of the brain. At the group level, individual statistical maps of learning trials were investigated in the context of a repeated-measures ANOVA to test the increase and decrease in brain activity during each learning condition, as well as differences between conditions. Additionally, we implemented conjunction analyses to test for brain regions that during learning commonly increased or decreased in BOLD signal across learning conditions, as expected from the previous findings (i.e. frontoparietal decrease and Default Mode Network increase), and that showed a learning condition-specific increase or decrease as compared to the other learning conditions. In the context of implementation trials, we implemented two separate group-level repeated-measures ANOVAs. One aimed to test for the repetition-wise linear increase and decrease per each learning condition and included a conjunction of contrasts to assess regions that showed a common increase/decrease across conditions. The other tested for differences between learning conditions as the mean across repetition levels and included a conjunction of contrasts to investigate differences in brain activity regarding the mean across repetition levels for each learning condition as compared to the others.

Based on the different hypotheses we meant to test for the across-condition and condition-specific signal change, respectively, we employed two different methodological approaches in the context of the conjunction analyses for learning and implementation trials alike. To elaborate, when testing for regions that exhibited a common increase or decrease across learning conditions in learning or implementation trials, we meant to reject the null hypothesis based on a number of activations k > 2, where 2 is the number of contrasts that are included in the conjunction (linear increase per each learning condition), i.e. 3 minus 1. On the other hand, when testing for brain regions where activity changed more for one condition than the others, either in a repetition-wise manner for learning trials or across-repetition in implementation trials, we used the global null hypothesis, which is based on a number of activations k = 0 (cf. congruent and incongruent contrasts, Friston et al., 2005; Nichols et al., 2005). Brain statistical maps inspection started at p(unc.) < .001. Results that are reported in this manuscript survived multiple comparison correction at the cluster level with p(FWE) < .05.

#### 2.6.3 Multivariate analysis

##### 2.6.3.1 Single trial estimation

In preparation for the identity-specific pattern similarity analysis, we estimated BOLD activity for individual trials using the Least Squares-Separate, LSS, approach (Mumford et al., 2012; Mumford et al., 2014). The analysis was conducted in SPM12 software package (Statistical Parametric Mapping; RRID:SCR_007037) and implemented in Matlab (version R2020b, The Math Works, Inc., 2020). For each experimental trial, correct and incorrect trials alike, we ran a single-trial GLM that included the event of interest, i.e. single trial, and all the other events, i.e. all the other trials plus the instruction cues, as two separate regressors. Event-related model regressors were constructed by modeling each event as a stick function convolved with the canonical HRF. The LSS analysis comprised a total of 768 single-trial GLMs (32 trials x 24 blocks) across 4 functional runs and included a high-pass filter at a 200s cut-off. The first 3 scans per run were discarded before starting the analysis.

##### 2.6.3.2 Identity-specific pattern similarity

The MVPA analysis was conducted using custom-made, in-house Matlab code, which incorporated functions from the CoSMoMVPA toolbox (version 1.1.0, Oosterhof et al., 2016) and its implementation in Matlab (version R2020b, The Math Works, Inc., 2020). The analysis included a whole-brain searchlight and an ROI-based MVPA. Single-trial activity T-maps were resampled from a 2 to a 3 isotropic voxel size before submitting them to both ROI-based and searchlight MVPA. This significantly reduced processing time. Importantly, different from the univariate analysis, in the MVPA we included correct and incorrect trials alike. The rationale for including incorrect trials was guided by the stimulus repetition structure, initially designed to prevent biased MVPA results (Ruge, Legler, et al., 2018). Excluding error trials would have disrupted the repetition chunk setup, as described in detail in section 2.2. In a similar vein, in the whole-brain as well as in the ROI-based MVPA only stimulus repetitions 2 to 8 were included. The exclusion of repetition 1 was motivated by the high error rate observed at this stimulus repetition level in the trial-and-error and observation-based learning conditions. Conversely, the error rate at stimulus repetition 2 in trial-and-error and observation-based learning trials was sufficiently low not to underestimate pattern similarities due to the inclusion of a disproportionate number of incorrect trials. Therefore, we coded the learning stage as stimulus repetitions 2 to 4, and the implementation stage as repetitions 5 to 8. For each learning block, rule-identity pattern similarities were estimated as the mean differences between correlation values for the re-occurrence of the same stimulus and the occurrence of different stimuli. Correlation values were based on vectors comprising activity values for the voxels within a searchlight sphere or within a predifined ROI. Pattern similiarities obtained for each block were averaged together for each learning condition.

##### 2.6.3.3 Searchlight

First, we implemented a spherical searchlight with a voxel, 9-mm radius, to produce a whole-brain map of regions where it was possible to decode rule-identities across all learning conditions and repetition levels. This was used to explore at a general level whether and where in the brain individual S-R rules could be decoded by means of all the information available we had in terms of learning conditions and stimulus repetitions (excluded stimulus repetition 1, see section 2.6.3.2). In addition, this whole-brain perspective also ensured that the more detailed pre-planned ROI-based MVPA would not miss relevant regions. At the group level, individual maps for stimulus repetitions 2 to 8, across learning conditions were submitted to one-sample t-tests and corrected for multiple comparisons at the cluster level in the context of FWE correction (p < .05).

##### 2.6.3.4 ROI-based MVPA

The ROI-based MVPA aimed at investigating where, within predefined ROIs, it was possible to decode rule identities. Going into more detail, as compared to the whole-brain searchlight analysis, the ROI-based analysis aimed at decoding rule identities for the re-occurrence of the same stimuli within and between stages (learning vs. implementation) and separately for the three different learning conditions (instruction, trial-and-error, and observation). In this way, we were able to compare the dynamics of rule representations from the learning stage to the implementation stage and to test whether rules could be better decoded in one stage with respect to the other, or similarly in both stages. Importantly, a significant pattern similarity effect that is common to both stages but tested separately per each stage does not imply that these similarity effects are based on the same representations. To test the consistency of rule representations between stages, we computed stimulus repetition pairwise comparisons from one stage to the other. Figure 2 depicts a schematic overview of how we implemented these analyses. In the *stage-specific analysis*, pattern similarity was separately determined within each stage so that each stimulus repetition was tested against each of the other stimulus repetitions within the stage of interest (Figure 2a). After the mean per each separate stage was computed, it was entered into a 3-way ANOVA with factors *stage* * *learning condition* * *ROI.* Importantly, a separate 3-way repeated-measures ANOVA was computed per each ROI group (Table 1). In the *cross-stage consistency analysis*, we assessed pattern similarity across stages by aggregating stimulus repetitions 2 to 8. This allowed us to identify stable rule identity patterns from the learning stage to the subsequent implementation stage. In this context, pattern correlation between repetitions within each stage were excluded so that each stimulus repetition of the learning stage was compared to each stimulus repetition of the implementation stage (Figure 2b). The mean of each learning-to-implementation pairwise comparison for the same stimulus repetition was then computed and compared to other stimulus repetitions learning-to-implementation pairwise comparison means in the context of a 2-way ANOVA with factors *learning condition* * *ROI.* A separate 2-way repeated-measures ANOVA was computed per each ROI group (Table 1).

**Figure 2.**
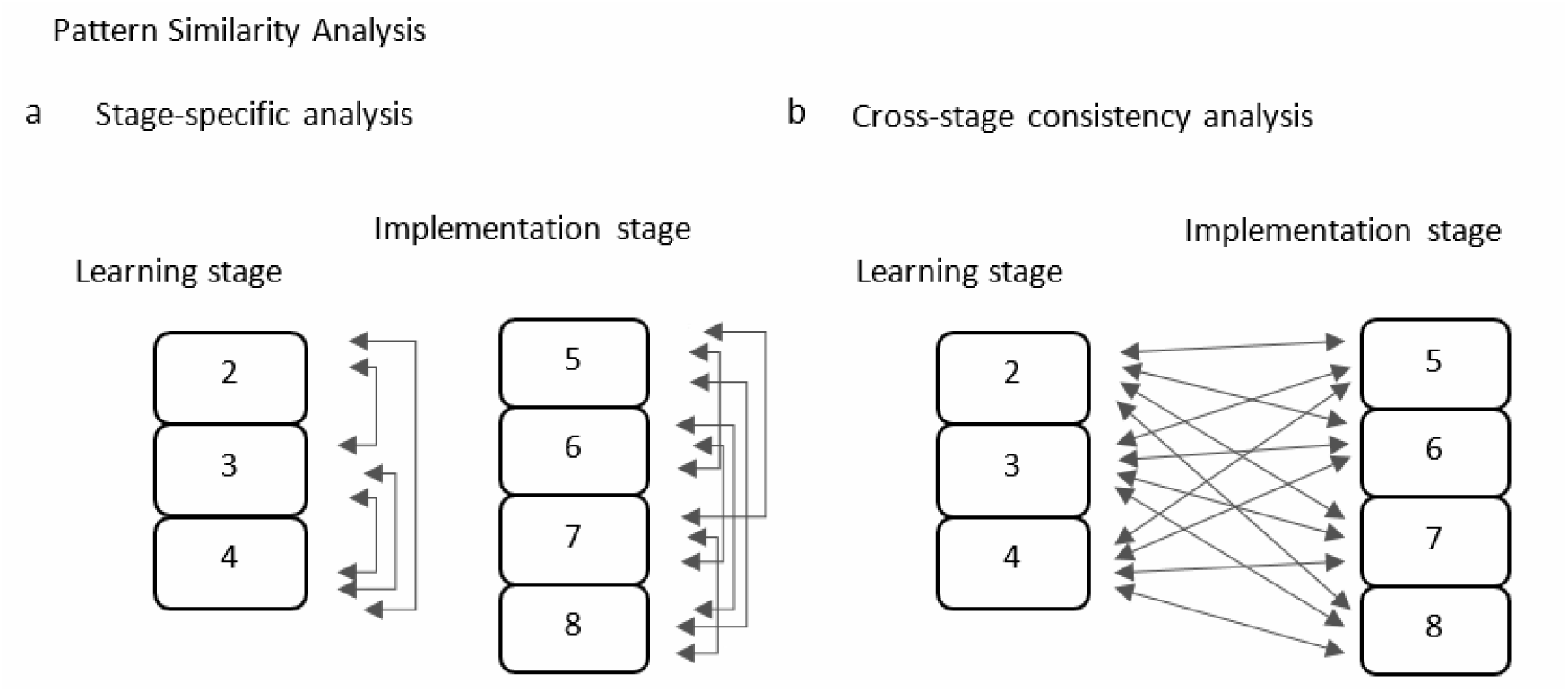
Schematic overview of pairwise comparisons in the pattern similarity analysis within the predefined ROIs. a) Stage-specific anaylsis: pattern similarities between stimulus repetitions were computed within each stage before averaging and comparing one stage to the other. b) Cross-stage consistency analysis: pattern similarity pairwise comparisons were computed between stages before averaging and comparing each mean value to the other.

**Table 1.** List of pre-planned ROIs for the MVPA.

| ROI_group | ROI | Hemisphere | Source |
| --- | --- | --- | --- |
| Prefrontal | VLPFC vs. DLPFC | L vs. R | AAL |
| Parietal | IPC vs. SPC | L vs. R | AAL |
| Premotor | vPMC/IFJ vs. dPMC | L vs. R | Searchlight across conditions and stages |
| Motor | Motor | L | Searchlight across conditions on implementation trials |

VLPFC = Ventrolateral Prefrontal Cortex; DLPFC = Dorsolateral Prefrontal Cortex; IPC = Inferior Parietal Cortex; SPC = Superior Parietal Cortex; vPMC = Ventral Premotor Cortex; IFJ = Inferior Frontal Junction; dPMC = Dorsal Premotor Cortex; Motor = Motor Cortex

###### Regions of interest

The predefined regions of interest included ventrolateral and dorsolateral PFC as in Ruge et al. (2019), as well as the superior and inferior parietal cortices in both hemispheres. Prefrontal and parietal ROIs were extracted from the AAL brain atlas that is available in the SPM12 toolbox WFU PickAtlas (Tzourio-Mazoyer et al. (2002); RRID:SCR_007378). The VLPFC ROI included the Inferior Frontal Gyrus (IFG) pars triangularis and opercularis, separately in the left and right hemispheres, whereas the DLPFC included the MFG, separately in the left and right hemispheres. In addition, we extracted ad-hoc ROIs starting from the whole-brain searchlight analysis. These were created via the SPM12 package bspmview (Spunt, 2016, v. 20180918). Specifically, we hypothesized that motor planning might play a role during learning across learning modes, even in the absence of overt motor response, like in instruction-based learning. This hypothesized role may be reflected in the brain by premotor rule decodability from the early stage of learning. Therefore, we extracted two 12-mm spherical ROIs in each hemisphere, in the ventral and dorsal premotor cortex. These were centered at different peak coordinates of the left and right premotor clusters that were found to contain S-R rule representations in the searchlight analysis that spanned all learning conditions and stimulus repetitions (right dPMC, MNI 42 -15 57; right vPMC/IFJ MNI 48 6 33; left dPMC -25 -13 63; left vPMC/IFJ MNI -43 3 42). Furthermore, based on the discovery that covert motor preparation preceding overt execution is reflected by effector-specific MVPA effects, as demonstrated by Ariani et al. (2022), we aimed to investigate whether the motor cortex contralateral to response implementation in implementation trials is involved during learning even when overt motor implementation is absent, as in the case of our instruction- and observation-based learning conditions. Hence, we extracted a 12-mm spherical ROI centered at the peak activation of the left motor cluster (MNI coordinates -46 -28 63) that exhibited decodable rules in the searchlight analysis across learning conditions on implementation trials only, stimulus repetitions 5, 6, 7 and 8. Since during implementation trials all learning conditions required response implementation, we ensured that the left motor cluster containing individual S-R rule representations could not be influenced by differences between conditions, as might have occurred during the learning stage. Our rationale was that if instruction-based learning involves motor planning via covert practice as early as during the instruction period, then this should be detectable in the pattern similarity analysis within the left motor ROI during instructed trials. Table 1 lists in detail the ROIs that we included in the subsequent MVPA.

Group-level statistics were implemented in RStudio (R Core Team, 2021). Single-subject pattern similarity values were submitted to a repeated-measures ANOVA (R package “afex,” Singmann et al., 2023) with within-subject factors learning condition and stage (plus brain region and hemisphere, where applicable (cf. Table 1) in the stage-specific analysis. For the cross-stage consistency analysis, the within-subject factor stage was not included in the model. Results were further investigated via post-hoc estimated marginal means tests (R package “emmeans,” Lenth, 2023). Reported results passed a significance threshold of p < .05 and were adjusted for the number of tests via the multivariate t-distribution approach.

## 3 Results

### 3.1 Behavior

#### 3.1.1 Trial-and-Error learning trials

A repeated-measures ANOVA on accuracy in trial-and-error learning trials revealed a main effect of stimulus repetition (F[2.68,212.00] = 1,255.02, MSE = 0.00455, *p* < .001, 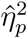 = .941) . A post-hoc analysis showed that accuracy in trial-and-error learning trials significantly increased from one stimulus repetition to the other (Figure 3a, all pairwise comparisons, t(79) > 7.5, p(t) < .0001).

**Figure 3.**
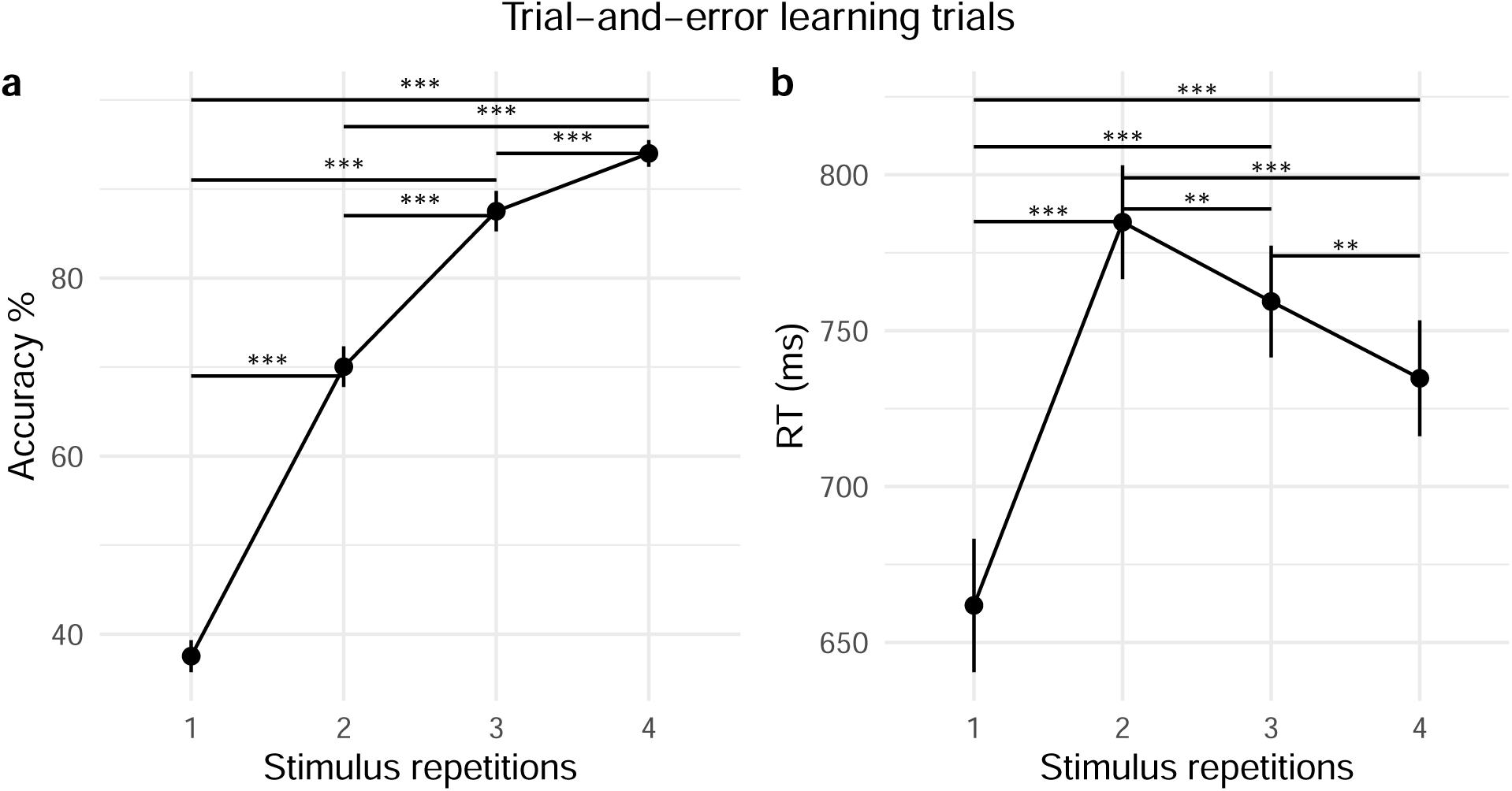
Behavioral performance increase per repetition level in Trial-and-Error learning trials. 95% confidence intervals are plotted.

A repeated-measures ANOVA on RT in trial-and-error learning revealed a main effect of stimulus repetition on RTs (F[2.01,159.04] = 63.42, MSE = 5,276.48, *p* < .001, 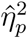 = .445). Post-hoc, RTs showed a significant increase from stimulus repetition 1, at which point participants were guessing, to stimulus repetition 2, during which they had to retrieve the previous S-R link. From stimulus repetitions 2 to 4, RTs significantly decreased from one repetition to the other (Figure 3b), probably reflecting task automatization. All pairwise comparisons between stimulus repetitions revealed to be significant (all comparisons, t(79) > 3.7, p(t) < .002).

#### 3.1.2 Implementation trials

For implementation trials, we computed a repeated-measures ANOVA on accuracy using within-subject factors *stimulus repetition* (1 to 4 in implementation trials) and *learning condition* (instruction, trial-and-error, observation). There was no effect of stimulus repetition on accuracy in implementation trials (F[2.84,224.73] = 1.40, MSE = 0.00081, *p* = .245, 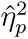 ) = .017, suggesting that learning successfully took place before the start of the implementation stage. Learning condition significantly affected accuracy during the implementation of previously learned S-R rules (main effect of learning condition, (F[1.87,147.50] = 5.89, MSE = 0.00281, *p* = .004, 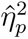 ) = .017. There was no interaction between the factors *stimulus repetition* and *learning condition* (F[5.03,397.13] = 2.07, MSE = 0.00093, *p* = .068, 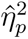 ) =.026 . A post-hoc analysis revealed a significantly greater accuracy in previously instructed as compared to previously observed trials (*t*(79) = 3.60, *p*_MV_*_t_*_(3)_ = .002; Figure 4a). No other pairwise comparisons between learning conditions reached significance.

**Figure 4.**
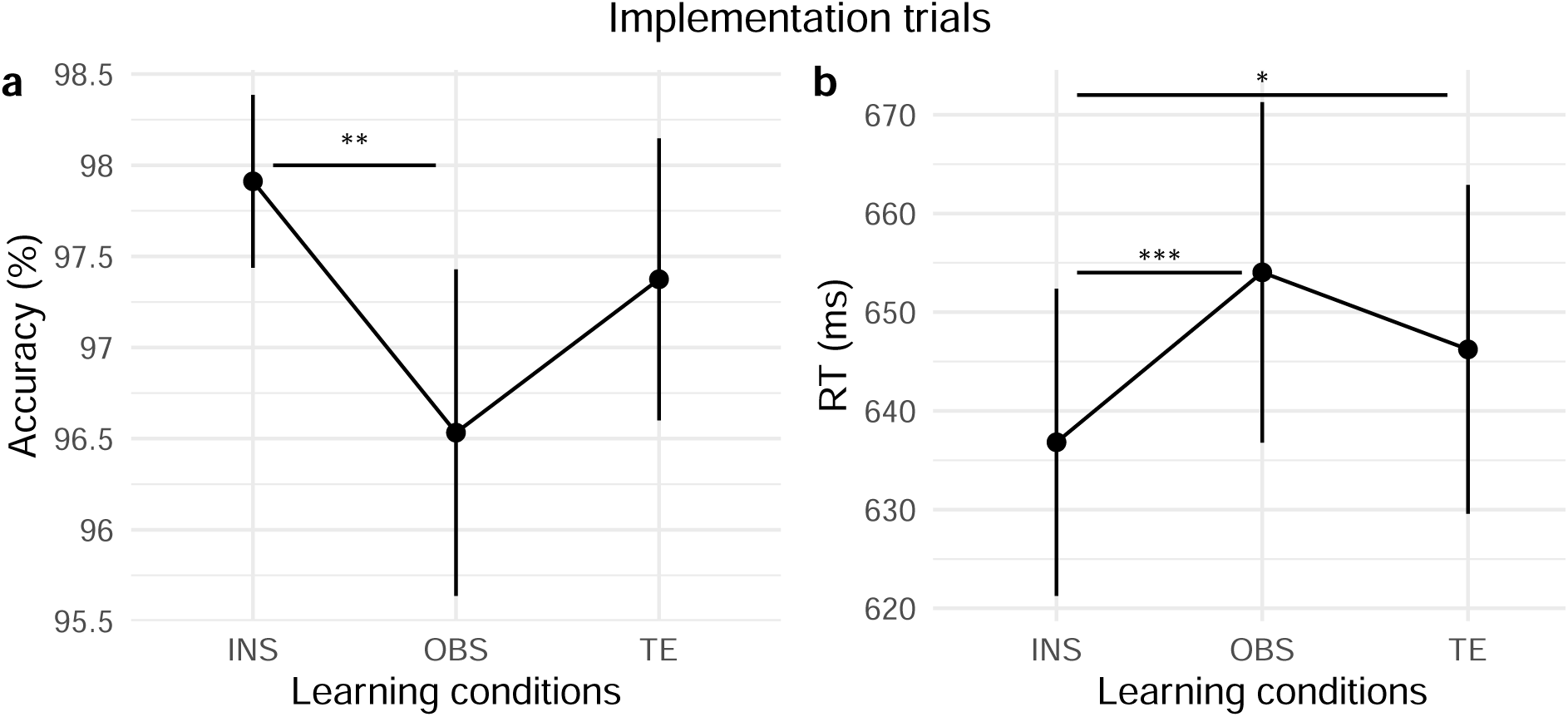
Difference between conditions in implementation trials. 95% confidence intervals are plotted.

The 2-way repeated-measures ANOVA to test the effect of *stimulus repetition* and *learning condition* on RT during implementation trials revealed that both, stimulus repetition (F[2.20,173.75] = 5.50, MSE = 1,752.56, *p* = .004, 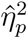 ) = .65) and learning condition (F[1.90,150.32] = 11.68, MSE = 2,137.52, *p* < .001, 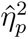 = .129) significantly affected RTs. There was no interaction effect between *stimulus repetition* and *learning condition* (F[5.06, 400.04] = 1.61, MSE = 973.59, *p* = .156, 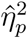 ). A post-hoc analysis showed that the effect of stimulus repetition was mainly guided by significantly faster RTs at the first stimulus repetition as compared to stimulus repetition 2 (*t*(79) = *−*3.50, *p*_MV_*_t_*_(6)_ = .004) and 3 (*t*(79) = *−*2.94, *p*_MV_*_t_*_(6)_ 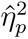 = .020). A post-hoc analysis testing for differences in RTs between learning conditions revealed a clear advantage of the instruction condition with faster RTs of previously instructed trials over both trial-and-error (*t*(79) = *−*2.85, *p*_MV_*_t_*_(3)_ = .015) and observation trials (*t*(79) = *−*5.03, *p*_MV_*_t_*_(3)_ *< .*001; Figure 4b).

### 3.2 Univariate fMRI analysis

#### 3.2.1 Common brain mechanisms during learning and implementation of S-R rules

Group-level statistical brain maps were first investigated in the context of repeated-measures ANOVAs testing for linear signal change during learning. In particular, we were interested in 1) common brain mechanisms underlying all learning modes and 2) signal change relative to a specific learning mode during learning.

##### 3.2.1.1 Learning stage

Figure 5 shows a statistical brain map of the regions that commonly increased (5a) and decreased (5b) in activity during learning across learning conditions. Results were investigated in the context of a conjunction analysis that included all learning conditions’ linear increase or decrease. Brain maps were investigated at p(unc.) < .001 and thresholded at the cluster level at p(FWE) < .05. The BOLD linear increase during learning across all three learning conditions included frontal, temporal, and parietal regions, as well as the bilateral precuneus, middle cingulate cortex, and left posterior cingulate cortex (cf. Table 2). Most of these regions have been linked to resting-state brain activity (for an overview, Raichle, 2015). It follows that their increased activity during learning across conditions is likely to mirror brain mechanisms related to decreasing cognitive control demand. In line with our predictions and the previous literature, prefrontal, parietal, and visual regions linearly decreased in their activity during learning (cf. Table 3). We interpreted this frontoparietal control network “relaxation” as the brain mechanism accompanying the aforementioned decreasing cognitive control demand that the DMN activation might hint towards. The decrease in occipital regions, on the other hand, likely resulted from mechanisms related to stimulus habituation due to repeated exposure during learning. In fact, in particular visuomotor learning seems to fine-tune those neurons that are responsive to the specific stimulus and response in occipital and motor regions (Makino et al., 2016 for a review on sensorimotor learning), which in turn would explain the decrease observed in these areas during learning.

**Figure 5.**
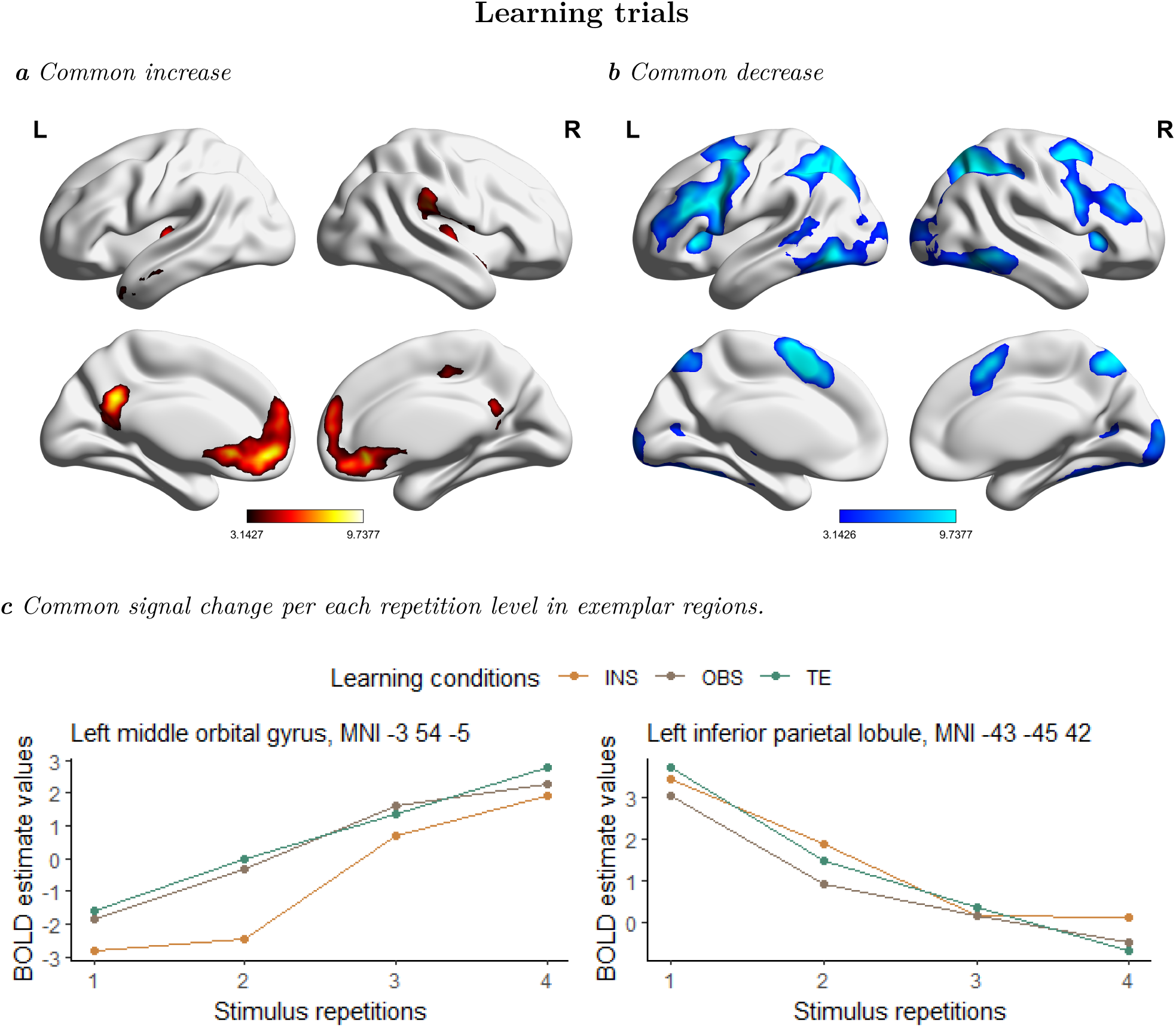
Across-condition signal change during S-R learning. a) Medial and lateral views of T-maps exhibiting increasing activity across learning conditions. b) Medial and lateral view of T-maps exhibiting decreasing activity across learning conditions. c) Stimulus-repetition-wise signal increase and decrease per each learning conditions (INS = instruction, OBS = observation, TE = trial-and-error) and stimulus repetition in exemplar brain regions. T-maps and BOLD estimate values result from the conjunction analysis (conjunction null) testing for brain regions with stimulus-repetition-wise signal increase or decrease common to all conditions during S-R learning. Results are FWE-cluster corrected (p < .05).

**Table 2.** Regions that linearly increased across learning conditions during learning.

| Region | MNI coordinates |  |  | Extent (voxels) | t-value (peak) |
| --- | --- | --- | --- | --- | --- |
|  | x | y | z |  |  |
| Frontal_Med_Orb_L | -3 | 54 | -5 | 3957 | 9.74 |
|  | -3 | 44 | -13 |  | 8.57 |
| Olfactory_L | -1 | 8 | -5 |  | 4.47 |
| ACC_sub_L | -3 | 24 | -7 |  | 7.83 |
| ACC_sub_R | 6 | 34 | -9 |  | 8.12 |
| ACC_pre_L | -13 | 48 | 10 |  | 5.75 |
| Frontal_Sup_Medial_L | -3 | 58 | 24 |  | 7.09 |
| Frontal_Sup_Medial_R | 6 | 58 | 12 |  | 6.81 |
| Frontal_Sup_2_L | -13 | 66 | 26 |  | 4.48 |
| Supramarginal_R | 52 | -31 | 30 | 2123 | 7.75 |
| Insula_R | 42 | -9 | -5 |  | 6.43 |
| Rolandic_Oper_R | 54 | 2 | 2 |  | 5.80 |
|  | 46 | -13 | 20 |  | 5.46 |
|  | 68 | -21 | 16 |  | 4.26 |
| Temporal_Sup_R | 66 | -5 | 2 |  | 4.70 |
|  | 42 | -31 | 16 |  | 3.99 |
| Insula_L | -39 | -17 | 6 | 1059 | 6.30 |
|  | -39 | 6 | -17 |  | 3.39 |
| Rolandic_Oper_L | -57 | -5 | 8 |  | 4.13 |
|  | -45 | -17 | 22 |  | 4.65 |
|  | -39 | -33 | 20 |  | 3.66 |
| Temporal_Sup_L | -41 | -7 | -9 |  | 5.32 |
|  | -67 | -15 | 4 |  | 4.25 |
| Temporal_Mid_L | -63 | -9 | -13 |  | 4.65 |
|  | -59 | 4 | -25 |  | 4.97 |
| Temporal_Pole_Mid_L | -49 | 14 | -31 |  | 4.63 |
| Supp_Motor_Area_R | 10 | -21 | 48 | 224 | 5.25 |

**Table 2***Regions that linearly increased across learning conditions during learning. (continued)*
| Region | x | y | z | Extent (voxels) | t-value (peak) |
| --- | --- | --- | --- | --- | --- |
| Cingulate_Mid_L | -7 | -19 | 46 |  | 3.21 |
| Cingulate_Post_L | -7 | -51 | 32 | 922 | 9.10 |
| Precuneus_L | -11 | -57 | 14 |  | 5.99 |
| Precuneus_R | 6 | -51 | 26 |  | 5.32 |

$T = 3.14$ , $p(\text{unc.}) < .001$ . Cluster-level $p(\text{FWE}) < .05$ ( $k = 224$ ). Regions were labeled using the AAL3 atlas. Peak separation threshold = 12 mm. Regions are organized anatomically by cluster, starting from the maximal t-value.

**Table 3.** Regions that linearly decreased across learning conditions during learning.

| Region | MNI coordinates |  |  | Extent (voxels) | t-value (peak) |
| --- | --- | --- | --- | --- | --- |
|  | x | y | z |  |  |
| Parietal_Inf_L | -43 | -45 | 42 | 26947 | 14.69 |
|  | -45 | -55 | 60 |  | 11.02 |
| Parietal_Inf_R | 50 | -39 | 50 |  | 10.35 |
| Angular_R | 30 | -53 | 44 |  | 13.41 |
| SupraMarginal_L | -59 | -53 | 32 |  | 4.54 |
| Parietal_Sup_L | -29 | -69 | 48 |  | 13.00 |
| Parietal_Sup_R | 18 | -71 | 58 |  | 10.52 |
| Precuneus_L | -9 | -61 | 54 |  | 6.63 |
| Temporal_Inf_L | -47 | -65 | -9 |  | 11.18 |
| Temporal_Inf_R | 54 | -55 | -11 |  | 9.79 |
| Temporal_Mid_L | -49 | -49 | 10 |  | 5.97 |
|  | -65 | -39 | -5 |  | 4.90 |
| Lingual_L | -25 | -85 | -17 |  | 5.67 |
| Fusiform_L | -37 | -23 | -25 |  | 4.65 |
| Fusiform_R | 36 | -41 | -23 |  | 6.45 |
| Occipital_Mid_R | 30 | -97 | 10 |  | 5.69 |

**Table 3***Regions that linearly decreased across learning conditions during learning. (continued)*
| Region | x | y | z | Extent (voxels) | t-value (peak) |
| --- | --- | --- | --- | --- | --- |
| Occipital_Sup_R | 34 | -73 | 42 |  | 12.06 |
| Calcarine_R | 10 | -97 | 8 |  | 7.12 |
| Calcarine_L | -11 | -103 | -5 |  | 6.84 |
| Cerebellum_6_R | 30 | -63 | -29 |  | 11.58 |
| Cerebellum_6_L | -31 | -61 | -29 |  | 10.78 |
| Cerebellum_Crus1_R | 50 | -77 | -21 |  | 4.68 |
| Cerebellum_Crus2_L | -7 | -77 | -37 |  | 6.39 |
| Cerebellum_8_R | 38 | -61 | -51 |  | 8.59 |
| Cerebellum_8_L | -35 | -67 | -53 |  | 7.49 |
| Cerebellum_10_L | -23 | -37 | -37 |  | 4.79 |
| Supp_Motor_Area_L | -7 | 14 | 52 | 18017 | 14.03 |
| Supp_Motor_Area_R | 6 | 8 | 76 |  | 3.87 |
| Cingulate_Mid_R | 12 | 16 | 36 |  | 5.72 |
| Precentral_L | -41 | 6 | 34 |  | 13.49 |
| Precentral_R | 48 | 10 | 32 |  | 9.48 |
| Frontal_Mid_2_L | -27 | -3 | 56 |  | 13.70 |
|  | -41 | 46 | 32 |  | 5.33 |
|  | -39 | 58 | 16 |  | 7.96 |
| Frontal_Mid_2_R | 44 | 40 | 32 |  | 10.39 |
| Frontal_Sup_2_R | 22 | 4 | 58 |  | 11.79 |
| Frontal_Inf_Tri_L | -41 | 28 | 22 |  | 12.13 |
| Frontal_Inf_Oper_R | 48 | 14 | 6 |  | 3.29 |
| Insula_L | -33 | 26 | -1 |  | 10.80 |
| Thal_PuA_L | -21 | -31 | 6 |  | 5.70 |
| Thal_VL_L | -13 | -13 | 10 |  | 9.58 |
| Insula_R | 34 | 22 | 2 | 369 | 9.12 |
| Thal_VL_R | 12 | -13 | 8 | 1336 | 7.71 |
| Thal_VPL_R | 26 | -25 | -1 |  | 4.72 |

**Table 3***Regions that linearly decreased across learning conditions during learning. (continued)*
| Region | x | y | z | Extent (voxels) | t-value (peak) |
| --- | --- | --- | --- | --- | --- |
| Putamen_R | 18 | 10 | 4 |  | 5.08 |
| Calcarine_R | 20 | -71 | 8 | 462 | 5.65 |
| Calcarine_L | -11 | -71 | 10 |  | 4.76 |
| Vermis_3 | -1 | -47 | -23 | 229 | 4.98 |

$T = 3.14$ , $p(\text{unc.}) < .001$ . Cluster-level $p(\text{FWE}) < .05$ ( $k = 229$ ). Regions labeled using the AAL3 atlas. Peak separation threshold = 20 mm. Regions are organized anatomically by cluster, starting from the maximal t-value.

Figure 5c shows two exemplar regions for the stimulus-repetition-wise linear increase and decrease in the left middle orbital gyrus, MNI -3 54 -5, and in the left inferior parietal lobule, MNI -43 -45 42, respectively.

##### 3.2.1.2 Implementation stage

Linear contrasts for the stimulus-repetition-wise signal increase and decrease per each learning condition were included in two different conjunction analyses testing brain regions that linearly increased and decreased, respectively, across learning conditions during the implementation of previously learned S-R rules.

Statistical maps were investigated at p(unc.) < .001 and corrected for multiple comparisons at the cluster level in the context of FWE-correction (p < .05). Figure 6 shows the statistical map of brain regions that linearly increased (6a) and decreased (6b) in activity during implementation trials across learning conditions. A linear increase was found in a bilateral occipital cluster, including the cuneus, motor regions in both hemispheres, as well as areas belonging to the DMN, like the angular gyrus, posterior cingulate, frontal and temporal cortex (for a complete list of activations, refer to Table 4). Brain regions in the left inferior parietal lobule, postcentral gyrus (Figure 6b), thalamus, and putamen linearly decreased in activity during S-R implementation trials, together with a right hemisphere cluster in the cerebellum (Table 5), possibly mirroring visual- and somatomotor adaptation as sensorimotor learning proceeds (e.g. Floyer-Lea & Matthews, 2004; Makino et al., 2016 for a review).

**Figure 6.**
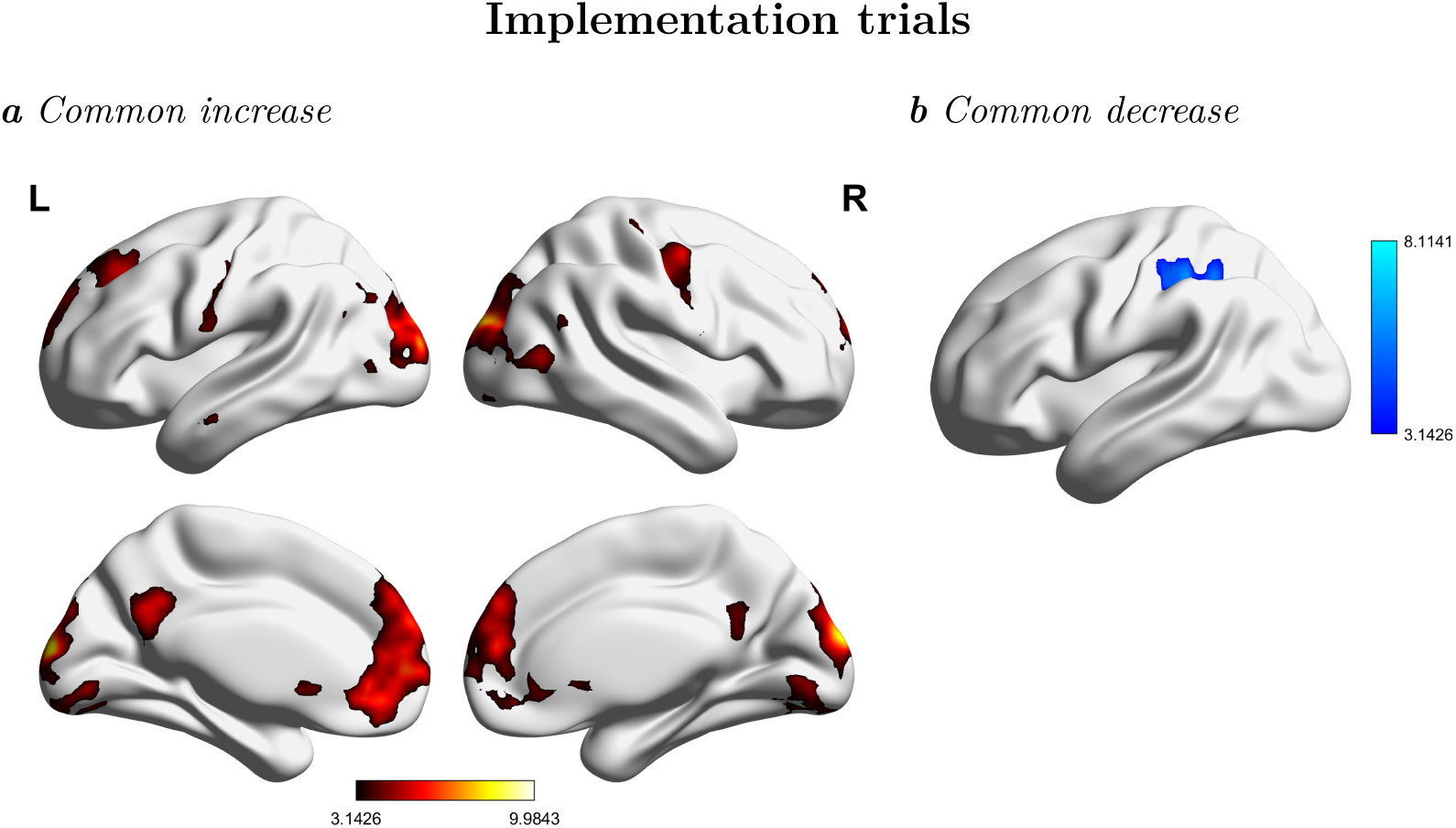
T-maps of the signal change across learning conditions in implementation trials. a) Medial and lateral view of brain regions exhibiting increasing activity across learning conditions. b) Lateral view of the left hemisphere (no suprathresholds cortical activations in the right hemisphere) with brain regions exhibiting decreasing activity across learning conditions. Statistical maps result from a conjunction analysis (conjunction null) testing for brain regions with stimulus-repetition-wise signal increase or decrease common to all conditions during S-R implementation. Results are FWE-cluster corrected (p < .05).

**Table 4.**
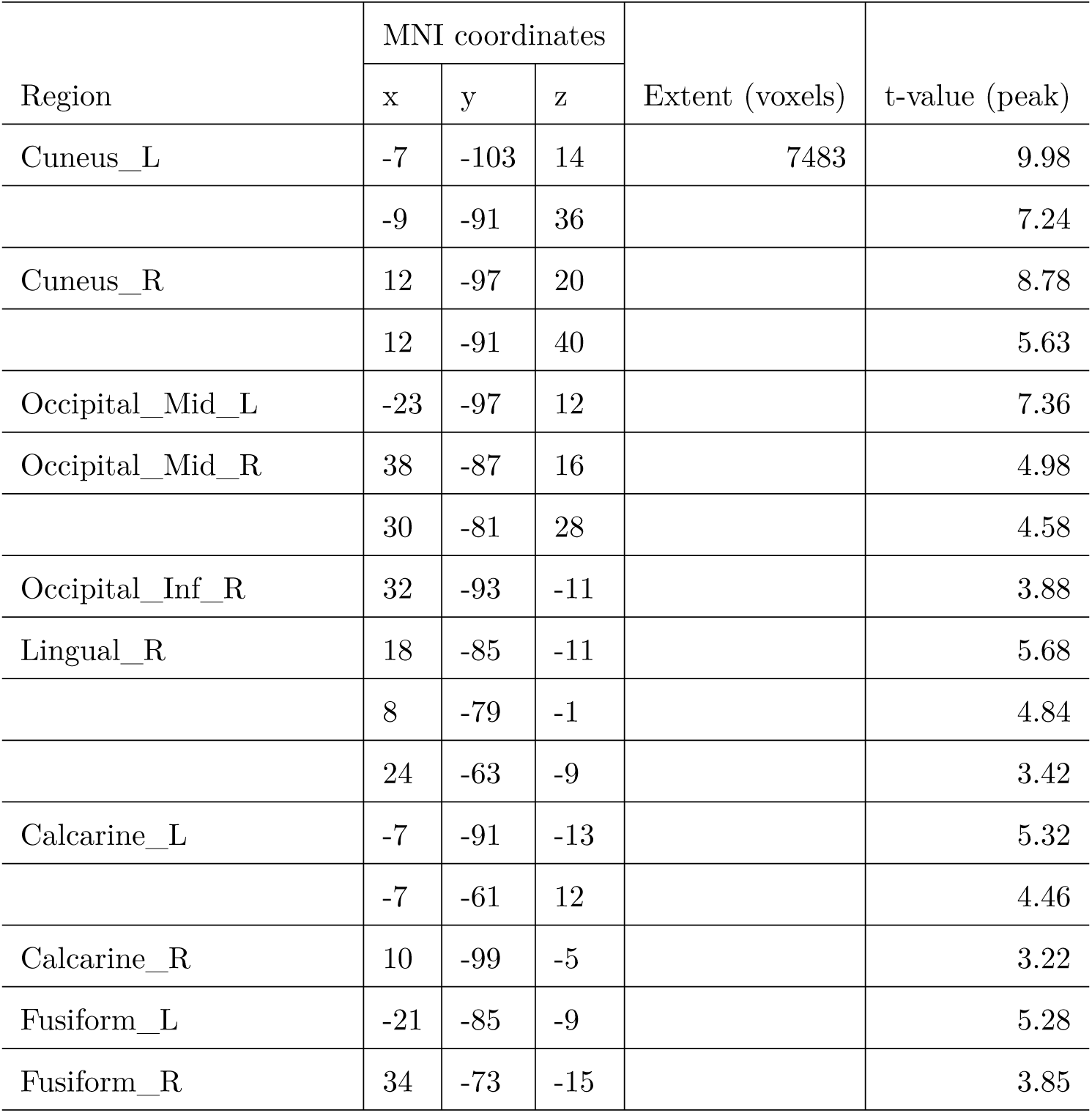
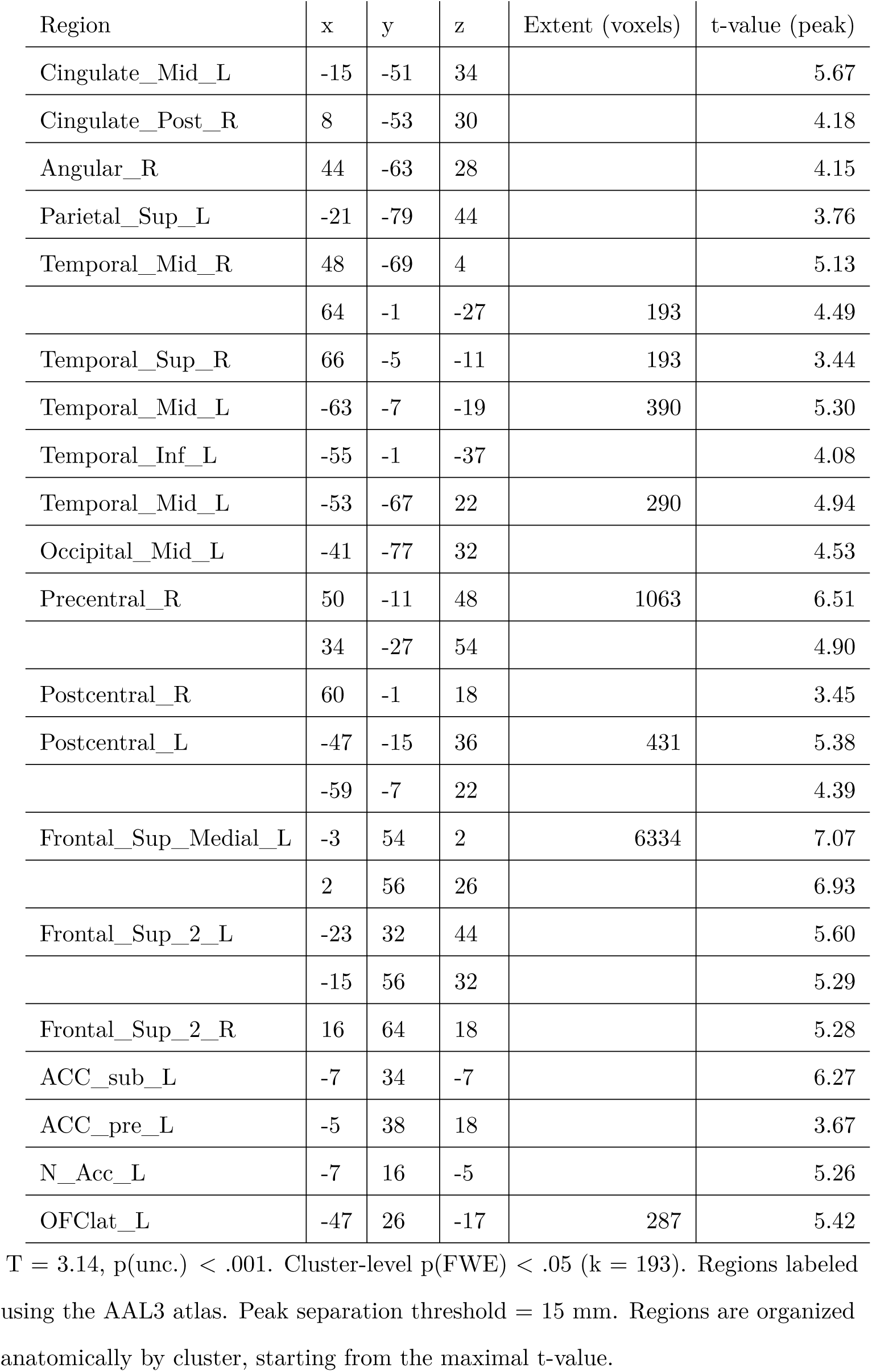
Regions that linearly increased across learning conditions during implementation trials.

| Region | MNI coordinates |  |  | Extent (voxels) | t-value (peak) |
| --- | --- | --- | --- | --- | --- |
|  | x | y | z |  |  |
| Cuneus_L | -7 | -103 | 14 | 7483 | 9.98 |
|  | -9 | -91 | 36 |  | 7.24 |
| Cuneus_R | 12 | -97 | 20 |  | 8.78 |
|  | 12 | -91 | 40 |  | 5.63 |
| Occipital_Mid_L | -23 | -97 | 12 |  | 7.36 |
| Occipital_Mid_R | 38 | -87 | 16 |  | 4.98 |
|  | 30 | -81 | 28 |  | 4.58 |
| Occipital_Inf_R | 32 | -93 | -11 |  | 3.88 |
| Lingual_R | 18 | -85 | -11 |  | 5.68 |
|  | 8 | -79 | -1 |  | 4.84 |
|  | 24 | -63 | -9 |  | 3.42 |
| Calcarine_L | -7 | -91 | -13 |  | 5.32 |
|  | -7 | -61 | 12 |  | 4.46 |
| Calcarine_R | 10 | -99 | -5 |  | 3.22 |
| Fusiform_L | -21 | -85 | -9 |  | 5.28 |
| Fusiform_R | 34 | -73 | -15 |  | 3.85 |

| Region | x | y | z | Extent (voxels) | t-value (peak) |
| --- | --- | --- | --- | --- | --- |
| Cingulate_Mid_L | -15 | -51 | 34 |  | 5.67 |
| Cingulate_Post_R | 8 | -53 | 30 |  | 4.18 |
| Angular_R | 44 | -63 | 28 |  | 4.15 |
| Parietal_Sup_L | -21 | -79 | 44 |  | 3.76 |
| Temporal_Mid_R | 48 | -69 | 4 |  | 5.13 |
|  | 64 | -1 | -27 | 193 | 4.49 |
| Temporal_Sup_R | 66 | -5 | -11 | 193 | 3.44 |
| Temporal_Mid_L | -63 | -7 | -19 | 390 | 5.30 |
| Temporal_Inf_L | -55 | -1 | -37 |  | 4.08 |
| Temporal_Mid_L | -53 | -67 | 22 | 290 | 4.94 |
| Occipital_Mid_L | -41 | -77 | 32 |  | 4.53 |
| Precentral_R | 50 | -11 | 48 | 1063 | 6.51 |
|  | 34 | -27 | 54 |  | 4.90 |
| Postcentral_R | 60 | -1 | 18 |  | 3.45 |
| Postcentral_L | -47 | -15 | 36 | 431 | 5.38 |
|  | -59 | -7 | 22 |  | 4.39 |
| Frontal_Sup_Medial_L | -3 | 54 | 2 | 6334 | 7.07 |
|  | 2 | 56 | 26 |  | 6.93 |
| Frontal_Sup_2_L | -23 | 32 | 44 |  | 5.60 |
|  | -15 | 56 | 32 |  | 5.29 |
| Frontal_Sup_2_R | 16 | 64 | 18 |  | 5.28 |
| ACC_sub_L | -7 | 34 | -7 |  | 6.27 |
| ACC_pre_L | -5 | 38 | 18 |  | 3.67 |
| N_Acc_L | -7 | 16 | -5 |  | 5.26 |
| OFClat_L | -47 | 26 | -17 | 287 | 5.42 |

$T = 3.14$ , $p(\text{unc.}) < .001$ . Cluster-level $p(\text{FWE}) < .05$ ( $k = 193$ ). Regions labeled using the AAL3 atlas. Peak separation threshold = 15 mm. Regions are organized anatomically by cluster, starting from the maximal t-value.

**Table 5.** Regions that commonly decreased across learning conditions during implementation trials.

| Region | MNI coordinates |  |  | Extent (voxels) | t-value (peak) |
| --- | --- | --- | --- | --- | --- |
|  | x | y | z |  |  |
| Postcentral_L | -45 | -35 | 50 | 1170 | 8.11 |
| Vermis_4_5 | -1 | -51 | -21 | 1389 | 6.98 |
| Vermis_8 | -1 | -63 | -35 |  | 6.36 |
| Cerebellum_6_R | 26 | -53 | -25 |  | 5.95 |
| Cerebellum_8_R | 18 | -61 | -47 |  | 5.11 |
| Thal_VL_L | -19 | -17 | 10 | 182 | 4.28 |
| Putamen_L | -25 | 2 | 12 |  | 4.21 |

$T = 3.14$ , $p(\text{unc.}) < .001$ . Cluster-level $p(\text{FWE}) < .05$ ( $k = 182$ ). Regions labeled using the AAL3 atlas. Peak separation threshold = 15 mm. Regions are organized anatomically by cluster, starting from the maximal t-value.

#### 3.2.2 Learning condition-specific signal change

##### 3.2.2.1 Learning stage

In order to find condition-specific brain activity during the learning stage, we performed conjunction analyses in the context of the repeated-measures ANOVA testing for a linear increase and decrease during learning. We were particularly interested in finding brain areas that linearly decreased or increased in activity significantly more in one learning condition than in the others. After p(FWE) cluster correction, clusters of regions only survived in the conjunction testing for regions that showed an increase specific to the instruction-based learning condition and a decrease specific to the trial-and-error learning condition (Figure 7a and 7b).

**Figure 7.**
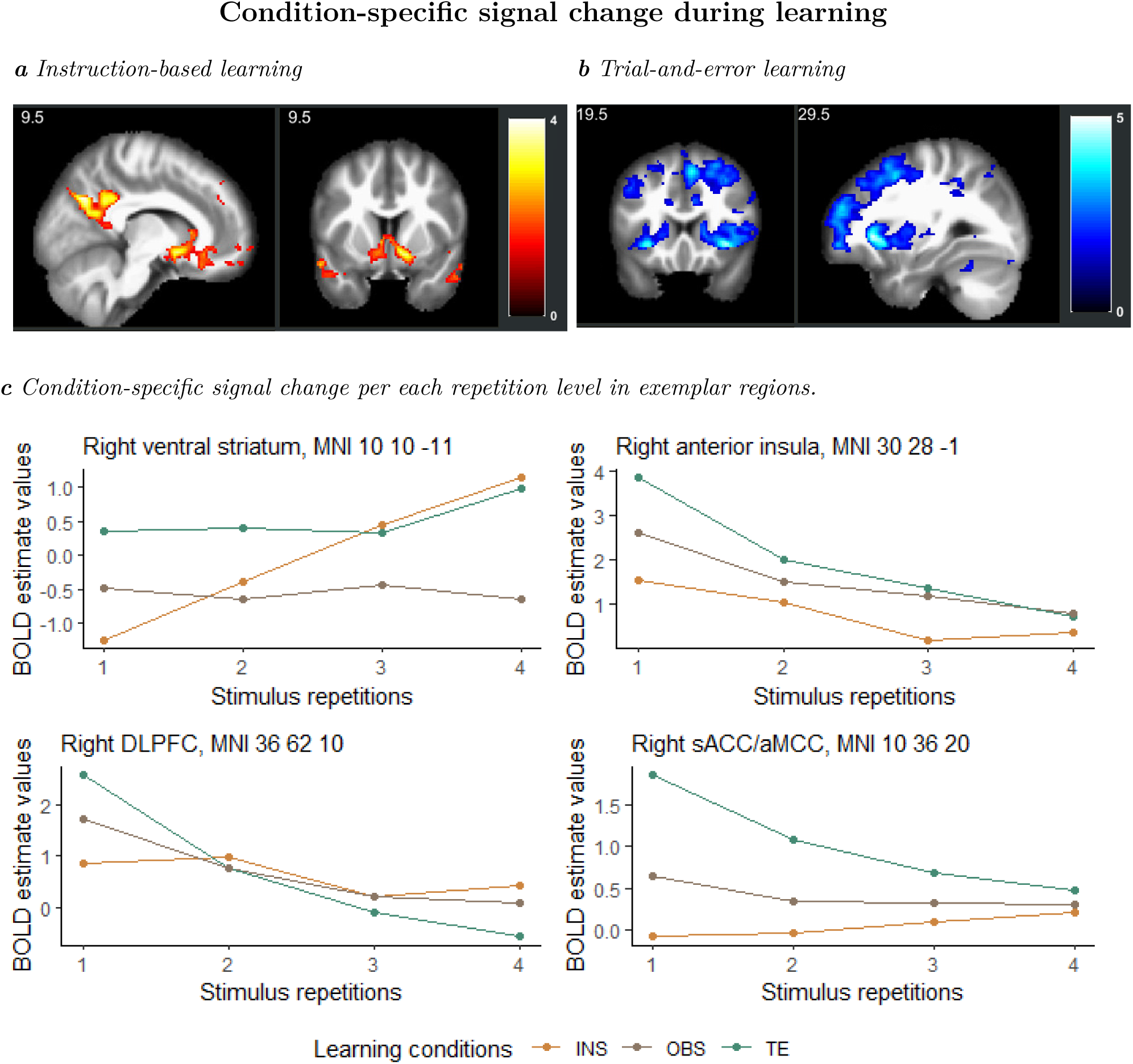
Condition-specific signal change during S-R learning. a) Coronal and sagittal view of the brain for instruction-specific linear increase as compared to trial-and-error and observed trials. b) Coronal and sagittal view of the brain for trial-and-error-specific linear decrease as compared to instructed and observed trials. c) Exemplar regions for condition-specific signal change. INS = instruction, OBS = observation, TE = trial-and-error, DLPFC = Dorso-Lateral Prefrontal Cortex. T-maps (a, b) and MNI coordinates (c) result from the conjunction analyses (global null, FWE-cluster corrected) that tested for condition-specific linear signal change during learning. BOLD estimate values come from repeated-measure ANOVA testing for the interaction learning condition * stimulus repetition.

###### Instruction-based learning-related signal change

Frontal, parietal and temporal areas commonly associated with the DMN showed a greater BOLD signal increase during instructed trials as compared to observed and trial-and-error trials (Figure 7a, Table 6). Additionally, regions in the basal ganglia showed a similar signal change pattern. Figure 7c shows the signal change in the right ventral striatum (i.e. Nucleus Accumbens, MNI 10 10 -11), as an exemplar region for the instruction-specific increase during learning.

**Table 6.**
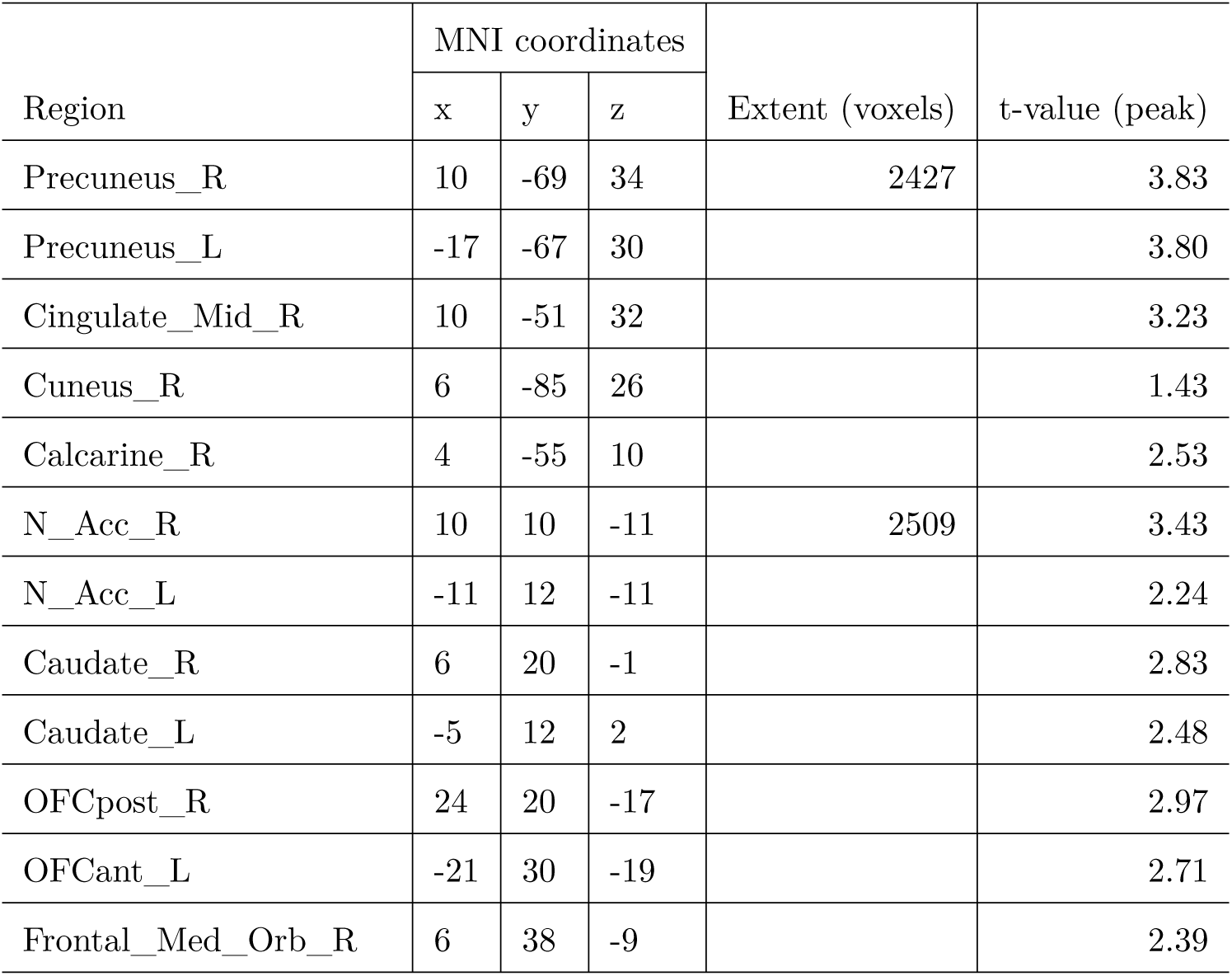
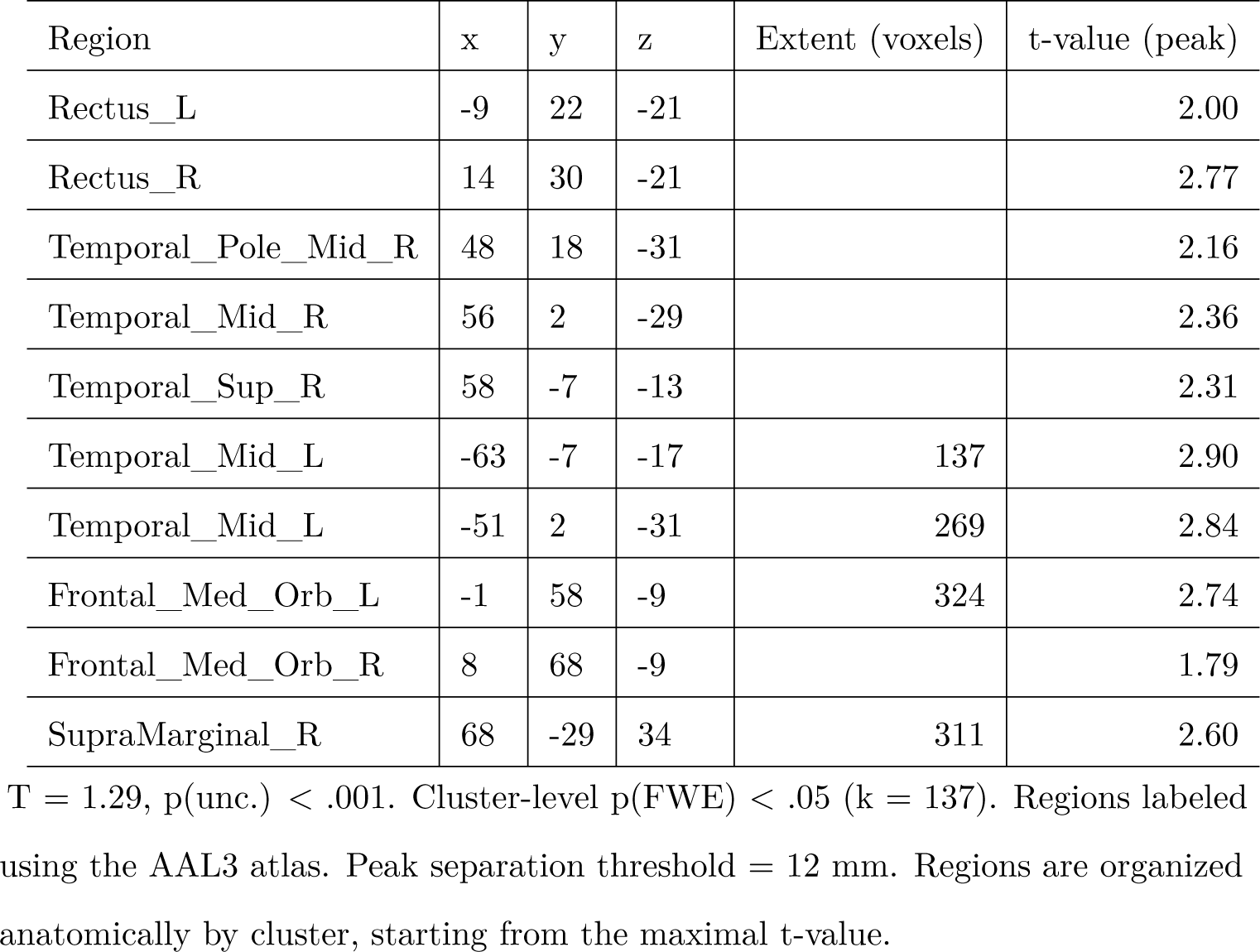
Regions with instruction-specific signal linear increase during learning.

###### Trial-and-error learning-related signal change

Frontal, prefrontal, and parietal regions, as well as bilateral clusters in the cerebellum (Figure 7b) linearly decreased in their activity more during trial-and-error learning than observed and instructed learning trials (Table 7). Figure 7c shows the signal change in the right anterior insula (AI, MNI coordinates 30 28 -1), right dorsolateral prefrontal cortex (DLPFC, MNI coordinates 36 62 10), and right supracallosal anterior cingulate cortex/anterior mid-cingulate cortex (sACC/aMCC, MNI coordinates 10 36 20), as exemplar regions for the trial-and-error-specific decrease as compared to instructed and observed trials.

**Table 7.** Regions with trial-and-error-specific signal decrease during learning.

| Region | MNI coordinates |  |  | Extent (voxels) | t-value (peak) |
| --- | --- | --- | --- | --- | --- |
|  | x | y | z |  |  |
| Supp_Motor_Area_R | 6 | 20 | 54 | 13249 | 3.98 |
| Frontal_Sup_Medial_L | -3 | 32 | 40 |  | 3.82 |
| ACC_sup_L | -13 | 36 | 22 |  | 2.69 |
| ACC_sup_R | 10 | 36 | 20 |  | 3.06 |
| Cingulate_Mid_R | 10 | 6 | 38 |  | 2.54 |
| Frontal_Sup_2_R | 36 | 62 | 10 |  | 3.85 |
| Frontal_Sup_2_L | -25 | 66 | 22 |  | 1.79 |
| OFCant_R | 24 | 46 | -13 |  | 3.46 |
| Frontal_Mid_2_L | -31 | 56 | -9 |  | 3.56 |
|  | -45 | 24 | 36 |  | 3.38 |

**Table 7***Regions with trial-and-error-specific signal decrease during learning. (continued)*
| Region | x | y | z | Extent (voxels) | t-value (peak) |
| --- | --- | --- | --- | --- | --- |
|  | -31 | 6 | 54 |  | 3.27 |
| Frontal_Mid_2_R | 42 | 36 | 36 |  | 3.01 |
|  | 30 | 10 | 50 |  | 3.46 |
| Frontal_Inf_Tri_R | 42 | 18 | 22 |  | 1.82 |
| Precentral_L | -31 | -13 | 64 |  | 2.84 |
| Precentral_R | 62 | 12 | 30 |  | 1.64 |
| Postcentral_L | -55 | -21 | 34 |  | 3.61 |
| Parietal_Inf_L | -47 | -59 | 50 |  | 3.23 |
| Supp_Motor_Area_L | -7 | -7 | 54 | 128 | 2.64 |
| Insula_R | 30 | 28 | -1 | 9732 | 4.78 |
| Insula_L | -31 | 22 | -7 |  | 4.18 |
| Frontal_Inf_Oper_R | 52 | 18 | 6 |  | 2.87 |
| Frontal_Inf_Oper_L | -49 | 6 | 6 |  | 1.73 |
| Putamen_L | -27 | -3 | -5 |  | 2.46 |
| Putamen_R | 30 | -3 | 4 |  | 2.28 |
| Caudate_L | -11 | 4 | 8 |  | 2.76 |
| Caudate_R | 12 | 10 | 16 |  | 3.23 |
| Thal_PuM_R | 4 | -27 | -1 |  | 3.62 |
| Thal_PuI_L | -19 | -25 | 12 |  | 2.26 |
| Thal_PuI_R | 16 | -21 | 16 |  | 2.06 |
| Cuneus_L | -7 | -77 | 18 |  | 2.63 |
| Lingual_R | 6 | -77 | -9 |  | 2.77 |
| Calcarine_L | 4 | -97 | -3 |  | 2.05 |
| Calcarine_R | 14 | -81 | 12 |  | 3.00 |
| Cerebellum_Crus1_L | -11 | -85 | -23 |  | 2.74 |
| Cerebellum_Crus1_R | 18 | -95 | -21 |  | 2.65 |
| Cerebellum_4_5_R | 18 | -55 | -17 |  | 3.11 |
| Cerebellum_6_L | -29 | -75 | -21 |  | 1.58 |

**Table 7***Regions with trial-and-error-specific signal decrease during learning. (continued)*
| Region | x | y | z | Extent (voxels) | t-value (peak) |
| --- | --- | --- | --- | --- | --- |
| SupraMarginal_R | 38 | -29 | 36 | 1691 | 2.50 |
| Angular_R | 38 | -61 | 38 |  | 2.43 |
| Parietal_Inf_R | 58 | -45 | 48 |  | 3.31 |
| Postcentral_R | 42 | -41 | 60 |  | 1.49 |
| Parietal_Sup_R | 30 | -71 | 58 |  | 1.43 |
| Precuneus_L | -7 | -69 | 42 | 1207 | 3.57 |
| Precuneus_R | 20 | -67 | 40 |  | 2.31 |

$T = 1.29$ , $p(\text{unc.}) < .001$ . Cluster-level $p(\text{FWE}) < .05$ ( $k = 128$ ). Regions labeled using the AAL3 atlas. Peak separation threshold = 20 mm. Regions are organized anatomically by cluster, starting from the maximal t-value.

##### 3.2.2.2 Implementation stage

Given the significant behavioral differences between conditions but not repetition levels in implementation trials, we computed group-level brain maps for the mean signal change across stimulus repetitions during implementation trials, separately for each condition. Across-repetition contrasts as the mean signal change per each learning condition were included in three conjunction analyses, each testing for signal change that was specific to one vs. the other conditions. However, no voxel survived correction, not even at p(unc.) < .001, in any of the conjunctions of contrasts, suggesting no main effect of learning condition at the brain level in implementation trials.

### 3.3 MVPA

#### 3.3.1 Searchlight

In order to find a whole-brain map to test whether and where individual S-R rules could be decoded in the brain in general and with the highest statistical power, we implemented a whole-brain spherical searchlight with a 3-voxel (9 mm) radius that included all learning conditions and repetition levels (excluding stimulus repetition 1). After cluster correction for multiple comparisons at p(FWE) < .05, the searchlight identified pattern similarity effect in extended bilateral clusters spanning occipital, prefrontal and parietal cortices (Figure 8, Table 8). Higher-order regions like prefrontal and parietal cortex likely reflect S-R rule representations, as opposed to occipital and motor cortices, which mirror stimulus and response identities, respectively (Ruge et al., 2019, p. Exp.2).

**Figure 8.**
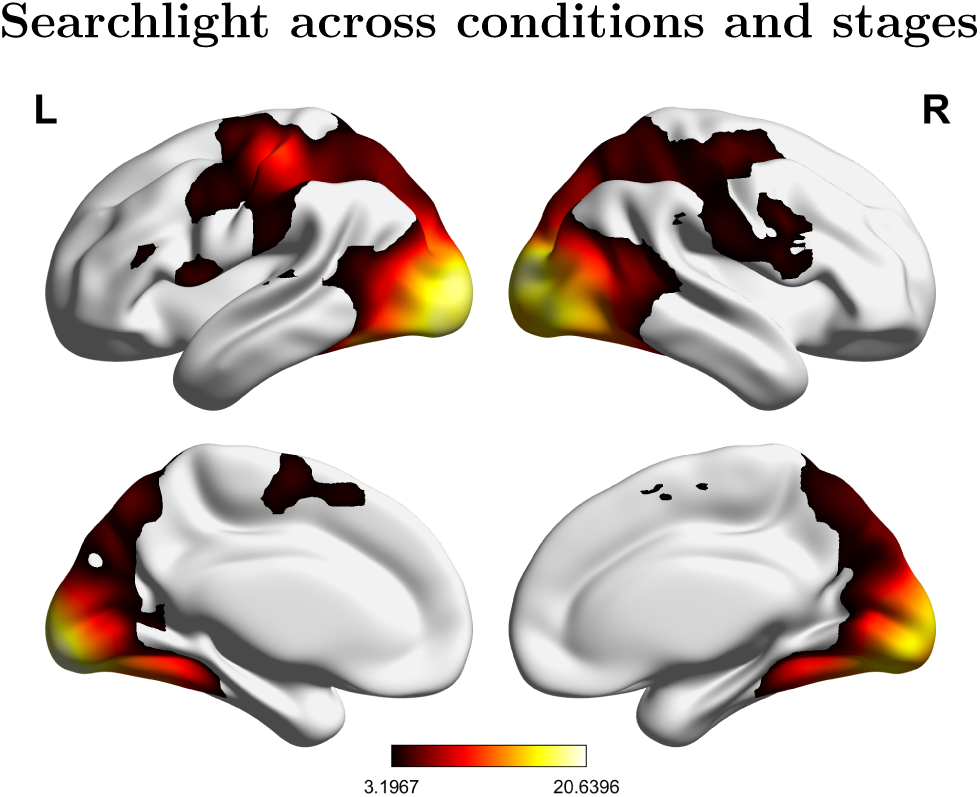
Whole-brain searchlight across learning conditions and repetition levels (stimulus repetitions 2 to 8). Statistical map was corrected at the cluster level in the context of FWE correction (p < .5, k = 19238).

**Table 8.** Whole-brain map of decodable S-R rules across conditions and stages.

| Region | MNI coordinates |  |  | Extent (voxels) | t-value (peak) |
| --- | --- | --- | --- | --- | --- |
|  | x | y | z |  |  |
| Occipital_Inf_L | -16 | -94 | -10 | 19238 | 20.64 |
| Occipital_Inf_R | 39 | -85 | -4 |  | 17.71 |
| Occipital_Sup_R | 27 | -97 | 12 |  | 19.18 |
| Occipital_Mid_L | -28 | -94 | 6 |  | 18.69 |
| Fusiform_L | -40 | -58 | -16 |  | 14.09 |
| Lingual_R | 21 | -88 | -7 |  | 17.37 |
| Temporal_Mid_L | -67 | -49 | 9 |  | 3.58 |
| Postcentral_L | -43 | -25 | 54 |  | 11.52 |
| Precentral_L | -25 | -13 | 63 |  | 5.89 |
|  | -43 | 3 | 39 |  | 5.28 |
| Precentral_R | 42 | -16 | 57 |  | 5.60 |
|  | 48 | 6 | 30 |  | 5.08 |
|  | 57 | 6 | 18 |  | 4.43 |

**Table 8** *Whole-brain map of decodable S-R rules across conditions and stages. (continued)*
| Region | x | y | z | Extent (voxels) | t-value (peak) |
| --- | --- | --- | --- | --- | --- |
| Supp_Motor_Area_L | -7 | -4 | 63 |  | 4.63 |
| Supp_Motor_Area_R | 3 | 6 | 57 |  | 4.50 |
| Paracentral_Lobule_L | -4 | -10 | 81 |  | 3.66 |
| Rolandic_Oper_R | 42 | -16 | 21 |  | 3.54 |
| Frontal_Inf_Tri_L | -49 | 30 | 18 |  | 4.10 |
| Frontal_Inf_Oper_L | -58 | 6 | 9 |  | 6.51 |
| Frontal_Sup_2_R | 27 | -1 | 54 |  | 4.82 |
| Insula_R | 36 | -7 | 21 |  | 3.60 |
| Precuneus_L | -1 | -70 | 54 |  | 6.01 |
| Parietal_Sup_L | -25 | -67 | 51 |  | 6.60 |
| Parietal_Sup_R | 30 | -58 | 57 |  | 5.68 |
| Parietal_Inf_R | 39 | -40 | 51 |  | 5.28 |
| SupraMarginal_L | -61 | -25 | 24 |  | 4.43 |
| SupraMarginal_R | 54 | -37 | 33 |  | 4.17 |

T = 3.20, $p(\text{unc.}) < .001$ . Cluster-level $p(\text{FWE}) < .05$ ( $k = 19238$ ). Regions labeled using the AAL3 atlas. Peak separation threshold = 12 mm. Regions are organized anatomically by cluster, starting from the maximal t-value.

#### 3.3.2 ROIs

##### 3.3.2.1 VLPFC and DLPFC

In the stage-specific analysis, where pattern similarities were determined separately within each stage, the repeated-measures ANOVA with within-subject factors *stage* * *condition* * *region* * *hemisphere* revealed an overall significant pattern similarity effect (Intercept, F[1,79] = 11.71, MSE = 0.00087, *p* < .001, 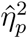 = .129). Pattern similarity values significantly differed between stages (main effect of stage, F[1,79] = 6.02, MSE = 0.00080, *p* = .016, 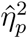 = .071) and hemispheres (main effect of hemisphere, F[1,79] = 9.65, MSE = 0.00007, *p* = .003, 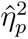 = .109), with learning > implementation (*t*(79) = 2.45, *p* = .016) and right > left (*t*(79) = *−*3.11, *p* = .003), respectively. Importantly, a significant pattern similarity effect was revealed only for the learning stage in both VLPFC and DLPFC (learning stage: VLPFC, *t*(79) = 3.41, *p* = .001, DLPFC *t*(79) = 3.04, *p* = .003; implementation stage: VLPFC *t*(79) = 1.62, *p* = .109, DLPFC, *t*(79) = 0.41, *p* = .683). In addition, stage and hemisphere significantly interacted (F[1,79] = 4.87, MSE = 0.00011, *p* = .030, 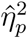 = .058), showing a more pronounced difference between stages in the right hemisphere (*t*(79) = 2.89, *p* = .005; Figure 9a) than in the left (*t*(79) = 1.64, *p* = .104).

**Figure 9.**
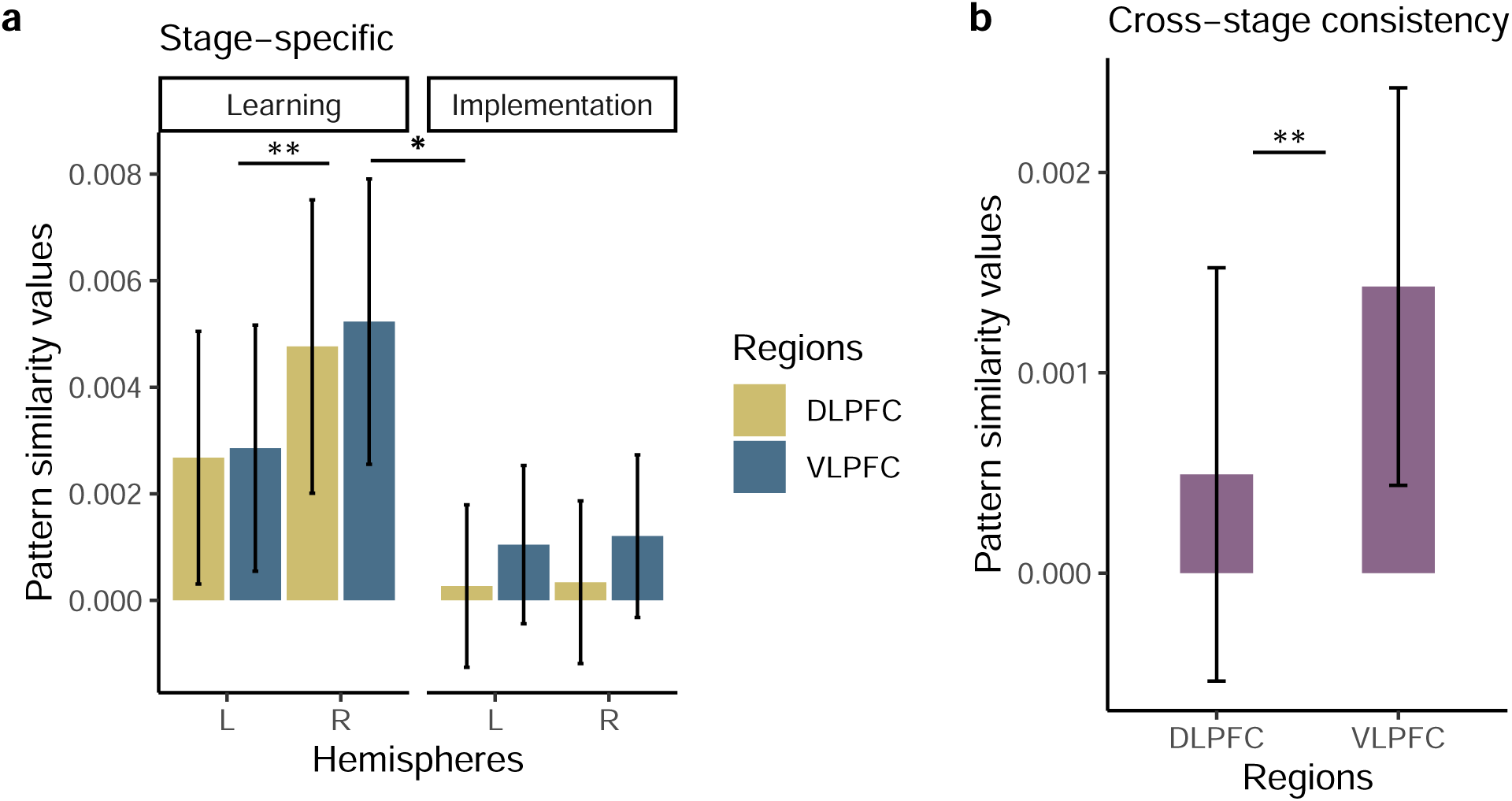
Pattern similarity effect in LPFC. 95% Confidence intervals are plotted.

When inspecting how consistent S-R rule representations were from the learning to the implementation stage, the repeated-measures ANOVA with within-subject factors *condition* * *region* * *hemisphere* revealed a marginal significant pattern similarity effect (Intercept, F[1,79] = 3.96, MSE = 0.00022, *p* .050, 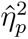 = .048). The cross-stage pattern analysis with within-subject factors *condition* * *region* * *hemisphere* showed a main effect of region (F[1,79] = 8.65, MSE = 0.00002, *p* = .004, 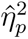 = .099) with a significant difference in rule identity patterns in VLPFC vs. DLPFC (*t*(79) = *−*2.94, *p* = .004; Figure 9b). There was no significant interaction with learning condition whatsoever, suggesting that S-R rules were represented similarly in the VLPFC across learning modes.

We speculated on these results given that the stage-specific pattern analysis revealed LPFC MVPA effects in learning but not implementation trials. We reasoned that the cross-stage consistency pattern analysis could be more sensitive than the stage-specific pattern analysis in picking up subtle but consistent patterns of activity in the implementation stage. The power of the cross-stage analysis in detecting consistency between representations, even those with subtler effects, is likely driven by the inclusion of a greater number of pairwise correlations spanning both stages (Figure 2).

##### 3.3.2.2 IPC and SPC

In the stage-specific MVPA within the IPC and SPC ROIs, the repeated-measures ANOVA with within-subject factors *stage* * *condition* * *region* * *hemisphere* revealed an overall significant pattern similarity significant effect (Intercept, F[1,79] = 43.27, MSE = 0.00089, *p* < .001, 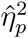 = .354). There was a significant impact of stage (F[1,79] = 6.93, MSE = 0.00092, *p* = .010, 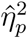 = .081) and region (F[1,79] = 8.50, MSE = 0.00014, *p* = .005, 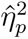 = .097) on S-R rule decodability, with learning > implementation (*t*(79) = 2.63, *p* = .010) and SPC > IPC (*t*(79) = *−*2.92, *p* = .005), as depicted in Figure 10a. In addition, stage and region significantly interacted with learning condition (F[1.83,144.29] = 3.53, MSE = 0.00011, *p* = .036, 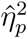 = .043) with significantly stronger pattern similarity values in the learning vs. implementation stage in the SPC during instructed trials (*t*(79) = 2.79, *p* = .007) and in the IPC during trial-and-error trials (*t*(79) = 2.35, *p* = .021).

**Figure 10.**
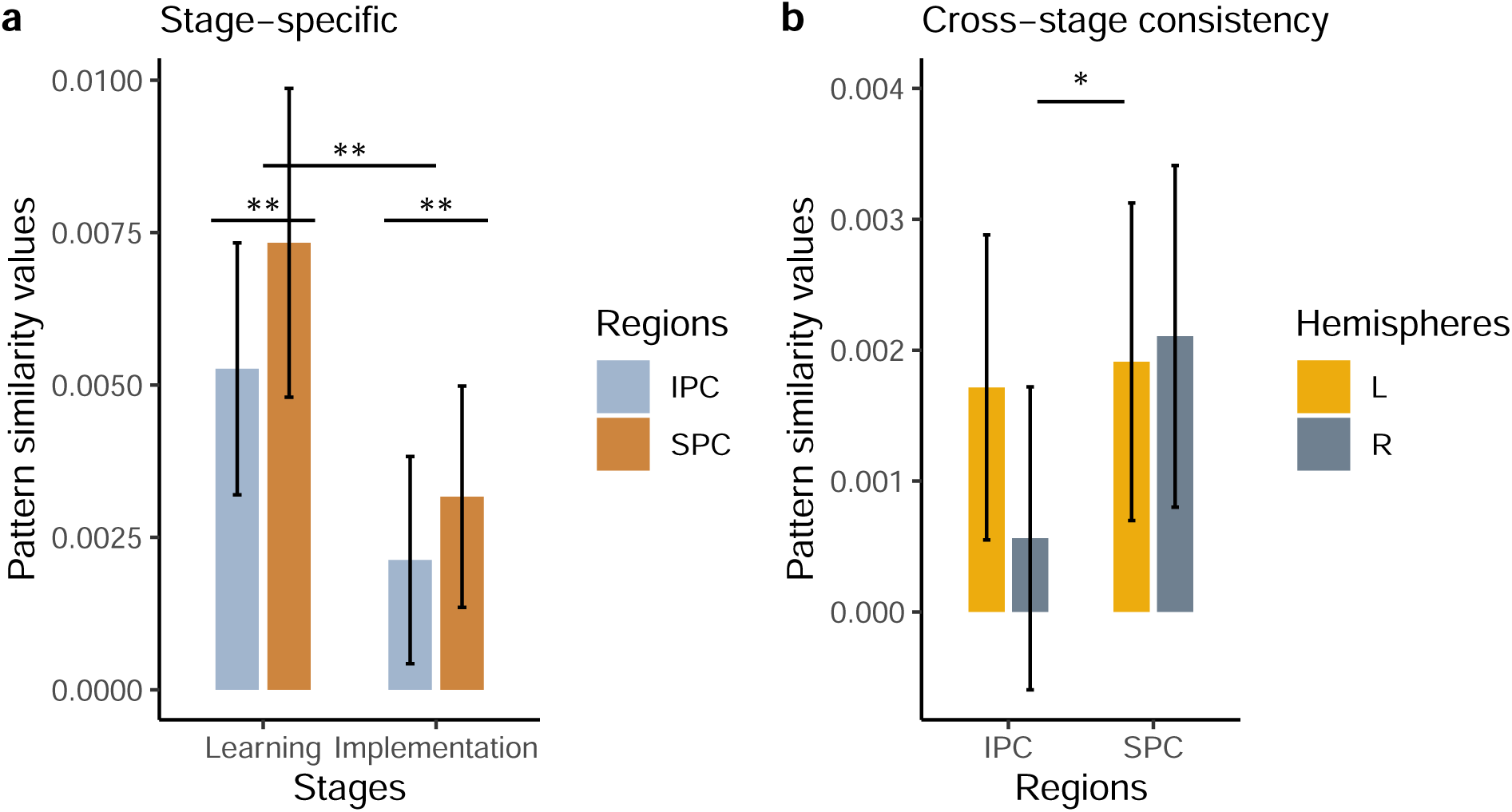
Pattern similarity effect in superior and inferior parietal cortex. 95% Confidence intervals are plotted.

The cross-stage consistency analysis with within-subject factors *condition* * *region* * *hemisphere* revealed again an overall pattern similarity significant effect (Intercept, F[1,79] = 9.14, MSE = 0.00026, *p* = .003, 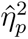 = .104). With a main effect of region (F[1,79] = 4.90, MSE = 0.00004, *p* = .030, 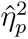 = .058), post-hoc estimated marginal mean comparison showed that S-R rules were represented consistently across stages more strongly in SPC vs. IPC (*t*(79) = *−*2.21, *p* = .030). Last, region and hemisphere significantly interacted (F[1,79] = 6.39, MSE = 0.00002, *p* = .013, 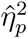 = .075). Post-hoc the interaction revealed to be mainly guided by higher pattern similarity values in the right SPC (*t*(79) = *−*3.04, *p* = .003; Figure 10b).

Overall, both prefrontal and parietal ROIs contained significantly decodable S-R rule representations, as suggested by the significant intercepts. Stronger pattern similarities were generally found in the learning vs. implementation stage and more in the right than in the left hemisphere. Notably, the VLPFC contained consistent cross-stage representations, in contrast with the DLPFC.

##### 3.3.2.3 vPMC/IFJ and dPMC

The stage-specific pattern analysis with within-subject factors *stage* * *condition* * *region* (ventral vs. dorsal PMC) * *hemisphere* revealed a significant overall similarity pattern effect (Intercept, F[1,79] = 37.82, MSE = 0.00064, *p* < .001, 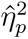 = .324). Particularly the right vPMC/IFJ appeared to contain significantly stronger pattern similarity values during learning as compared to implementation trials (3-way interaction *stage* * *region* * *hemisphere*, F[1,79] = 4.10, MSE = 0.00011, *p* = .046, 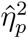 = .049, Figure 11a). Importantly, individual S-R rule representations were significantly apparent across stages, regions, and hemispheres as suggested by testing the individual marginal means for the 3-way interaction *stage* * *region* * *hemisphere* against 0 (all individual interactions per each *stage* * *region* * *hemisphere* combination, t[79] > 2.39, p < .019).

**Figure 11.**
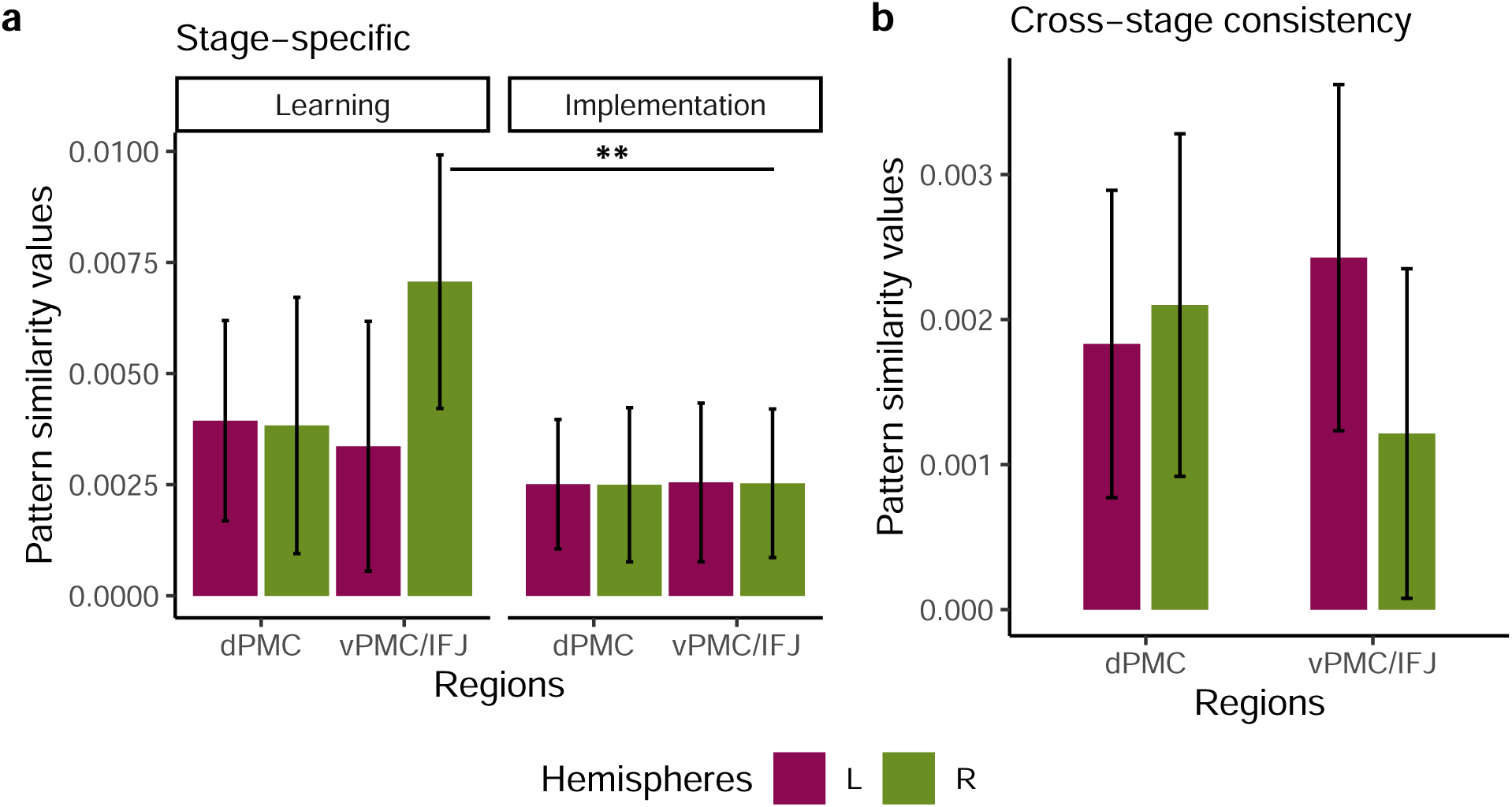
Pattern similarity effect in Premotor Cortex, PMC. dPMC = Dorsal Premotor Cortex; vPMC/IFJ = Ventral Premotor Cortex/Inferior Frontal Junction. a) Stage-specific analysis. b) Cross-stage consistency analysis. 95% Confidence intervals are plotted.

A general pattern similarity effect was apparent from the cross-stage consistency analysis with within-subject factors *condition* * *region* * *hemisphere* (Intercept, F[1,79] = 21.63, MSE = 0.00016, *p* < .001, 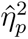 = .215), suggesting that stable cross-stage representations of individual S-R rules were contained within all of the premotor ROIs (Figure 11b). The analysis did not show any main effect of condition or stage, nor a significant interaction.

##### 3.3.2.4 Left Motor ROI

The group-level repeated-measures ANOVA with within-subject factors *condition* and *stage* revealed an overall significant pattern similarity effect in the left motor ROI (Intercept, F[1,79] = 105.92, MSE = 0.00055, *p* < .001, 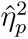 = .573). The ANOVA model additionally showed a significant effect of condition (F[1.89,149.02] = 4.69, MSE = 0.00040, *p* = .012, 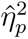 = .056) with significantly stronger pattern similarities in trial-and-error as compared to observed trials (*t*(79) = 1.73, *p*_MV_*_t_*_(3)_ = .198, *t*(79) = *−*1.41, *p*_MV_*_t_*_(3)_ = .337, *t*(79) = *−*2.84, *p*_MV_*_t_*_(3)_ = .015). Importantly, there was no interaction between stage and condition (F[1.97,155.72] = 1.00, MSE = 0.00050, *p* = .368, 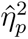 = .013) as one could have expected from the response requirement in trial-and-error learning but not in instructed and observed learning trials, nor a significant effect of stage on motor representations pattern similarity (F[1,79] = 1.45, MSE = 0.00053, *p* = .233, 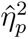 = .018). In fact, one could have predicted the implementation stage, which involved overt motor responses in all learning conditions, to yield significantly greater pattern similarity values compared to the learning stage, where learning was based on the passive view of S-R rules in the instruction- and observation-based learning conditions. Therefore, we inspected motor representations in the learning and implementation stages, for each condition separately. Pattern similarity values were significantly different from 0 across learning conditions in the implementation stage (trial-and-error, *t*(79) = 6.59, *p*_MV_*_t_*_(6)_ *< .*001; instruction, *t*(79) = 7.74, *p*_MV_*_t_*_(6)_ *< .*001; observation, *t*(79) = 5.51, *p*_MV_*_t_*_(6)_ *< .*001), as expected from the common motor response implementation across conditions. However, only trial-and-error and instructed but not observed trials led to pattern similarities values significantly different from 0 in the learning stage (trial-and-error, *t*(79) = 5.53, *p*_MV_*_t_*_(6)_ *< .*001; instruction, *t*(79) = 3.39, *p*_MV_*_t_*_(6)_ = .006; observation, *t*(79) = 1.63, *p*_MV_*_t_*_(6)_ = .476; Figure 12a). When testing for differences between learning conditions separately for each stage, there was a trend towards significance for trial-and-error > observation in learning trials (*t*(79) = *−*2.33, *p*_MV_*_t_*_(3)_ = .057), which, we reasoned, could have been the factor driving the main effect of condition that we initially found (and that referred to pattern similarity values averaged over the levels of stage, i.e. learning and implementation). Importantly, omitting correction for multiple comparisons led to a statistically significant difference between trial-and-error and observation in learning trials (*t*(79) = *−*2.33, *p* = .022). Meanwhile, the comparison between instruction and trial-and-error learning trials still yielded no significant difference (*t*(79) = *−*1.50, *p* = .138), confirming the presence of decodable response information in motor cortex in instruction-based learning trials.

**Figure 12.**
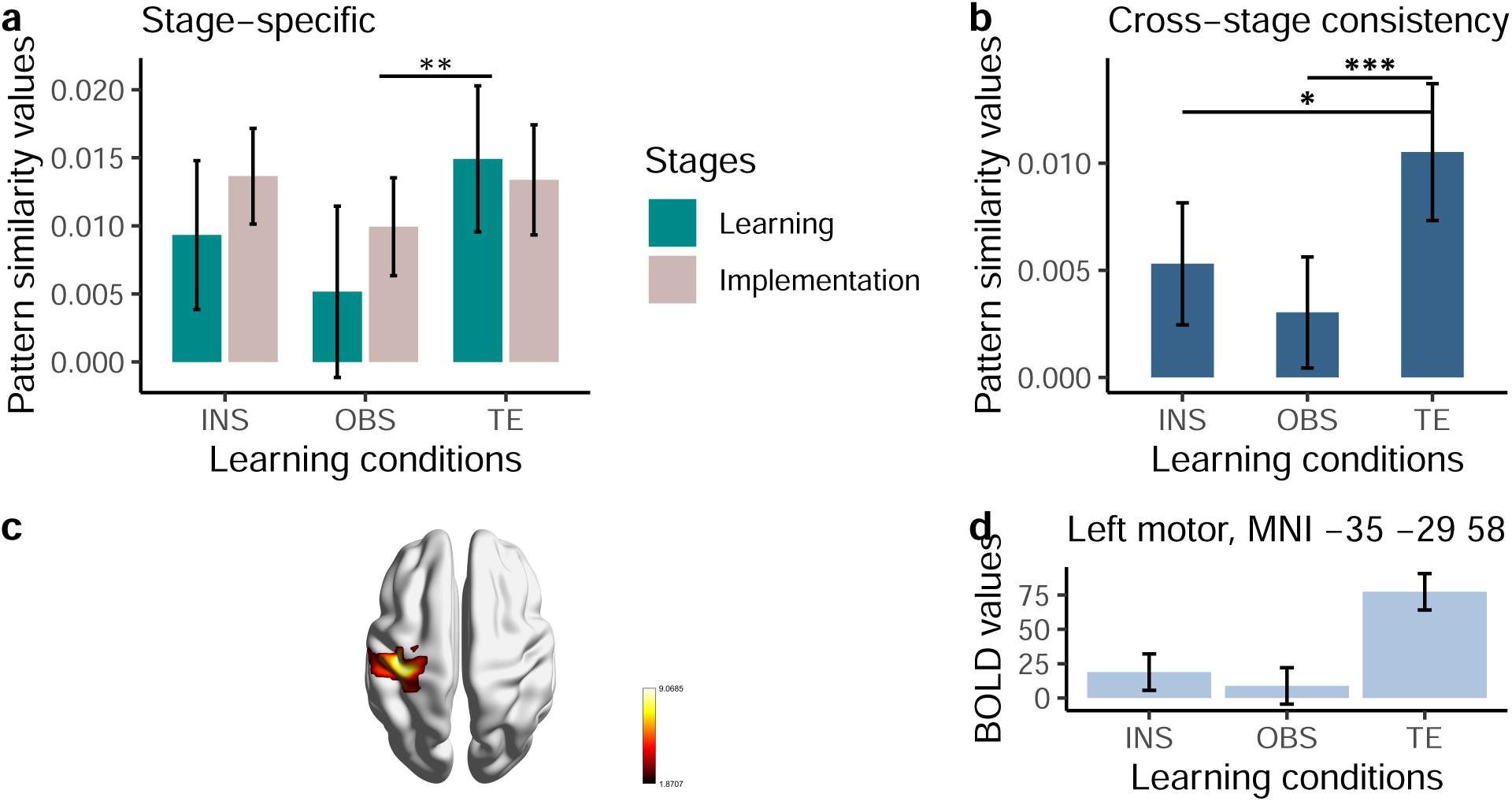
Pattern similarity effect in Left Motor ROI in the a) Stage-specific analysis, and b) Cross-stage consistency analysis. INS = Instruction-based learning; OBS = Observation-based learning; TE = Trial-and-error learning. c) Left motor cluster from conjunction analysis (FWE peak-level correction) for regions significantly active in trial-and-error > instruction- and observation-based learning. d) BOLD estimates plotted at the global maximum MNI coordinates [MNI -35 -29 58] of the left motor cluster as in c). 95% Confidence intervals are plotted.

To sum up, the additional investigation of stage-wise pattern similarity effects within the left motor ROI showed that already during learning motor responses were represented in the absence of motor program enactment in instructed but not observed, as in our observation-based learning condition, trials. Additionally, motor responses were represented during learning in the same regions that contained pattern similarities during S-R rule implementation. This, we suggest, may reflect motor planning in the form of covert execution and motor imagery.

The cross-stage consistency analysis with within-subject factor *condition* showed an overall significant representational stability (F[1,79] = 46.79, MSE = 0.00020, *p* < .001, 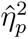 = .372), with consistent cross-stage motor representations in instruction and trial-and-error, but not observation trials (t-test against 0 for instruction, *t*(79) = 3.71, *p*_MV_*_t_*_(3)_ = .001; trial-and-error, *t*(79) = 6.56, *p*_MV_*_t_*_(3)_ *< .*001; observation, *t*(79) = 2.33, *p*_MV_*_t_*_(3)_ = .065). The repeated-measures ANOVA additionally revealed a main effect of condition (F[1.94,153.27] = 7.78, MSE = 0.00016, *p* = < .001, 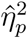 = .090), which post-hoc unfolded a significantly stronger cross-stage consistent decodability in trial-and-error as compared to both instruction- (*t*(79) = *−*2.48, *p*_MV_*_t_*_(3)_ = .040) and observation-based learning conditions (*t*(79) = *−*3.92, *p*_MV_*_t_*_(3)_ *< .*001; Figure 12b).

###### Follow-up univariate analysis of motor activity

Despite the indisputable advantage of MVPA as compared to the standard GLM in detecting patterns of activity related to a certain cognitive process, we were interested in testing whether the rule representation-related left motor activation that was apparent from the MVPA was also detectable in the standard univariate analysis of learning trials. Our working hypothesis was that mere motor preparation in the absence of overt motor implementation, as apparent from the multivariate pattern similarity analysis in instruction-based learning during the learning stage, would be associated with drastically weaker activity in the univariate analysis as compared to trial-and-error learning trials, which required motor implementation. This hypothesis was additionally guided by Ariani et al.’s results (2022) in showing that motor preparation was associated with effector-specific MVPA but not univariate effects in the motor cortex. Therefore we were interested in finding a left motor activity cluster that we could make sure it was related to response preparation, like in the trial-and-error learning condition, and in testing whether this was significantly activated during instructed and observed learning trials, which did not require response implementation during learning.

We implemented (mean) across-repetition t-contrasts for differences between conditions and a conjunction analysis testing for brain regions with increased BOLD signal in trial-and-error vs. instruction and trial-and-error vs. observation (global null for congruent contrasts) during the learning stage. Reported tables and figures show results after FWE peak level correction. We corrected at the peak and not cluster level for cluster-specific visualization purposes. The analysis showed a significant left motor cluster with global maximum at MNI -35 -29 58 (Figure 12c; Table 9). BOLD estimate values at the global maximum peak coordinates, showed, in line with our hypothesis, a very weak motor activation during learning when response implementation was not required as compared to trial-and-error learning (Figure 12d). Similar results were observed at distinct MNI coordinates within the left motor cluster, both from the whole-brain searchlight analysis on implementation trials and the brain map depicting heightened activity during S-R rule motor execution across repetition levels and learning conditions.

**Table 9.** Table of the left motor cluster activation from the conjunction analysis testing for regions significantly actuve in trial-and-error > instruction- and observation-based learning.

| Region | MNI coordinates |  |  | Extent (voxels) | t-value (peak) |
| --- | --- | --- | --- | --- | --- |
|  | x | y | z |  |  |
| Precentral_L | -35 | -29 | 58 | 5061 | 9.07 |
|  | -25 | -19 | 78 |  | 6.00 |
| Rolandic_Oper_L | -49 | -23 | 20 |  | 8.29 |
| Postcentral_L | -63 | -15 | 32 |  | 4.38 |
| Parietal_Inf_L | -57 | -23 | 50 |  | 8.62 |

**Table 9***Table of the left motor cluster activation from the conjuncti (continued)*
| Region | x | y | z | Extent (voxels) | t-value (peak) |
| --- | --- | --- | --- | --- | --- |
| Putamen_L | -23 | 6 | -5 |  | 4.60 |
| Insula_L | -39 | -3 | 14 |  | 7.55 |

T = 3.27, peak-level FWE-correction ( $p < .05$ ). Regions labeled using the AAL3 atlas. Peak separation threshold = 20 mm. Regions are organized anatomically by cluster, starting from the maximal t-value.

Together these results suggest that covert motor preparation in the absence of overt motor implementation during learning, as in our instruction- and observation-based learning conditions, is associated with drastically weaker left motor activity in the univariate analysis as compared to trial-and-error trials, which required motor implementation during learning. This contrasts with the multivariate pattern analysis, which showed statistically similar rule decodability in instruction-based and trial-and-error learning during the learning stage. Our results are in line with the recent work by Ariani et al. (2022), who showed the advantage of the multivariate approach in unfolding covert motor preparation in the absence of motor implementation. in turn, this seems to provide firm evidence for motor preparation, i.e. proceduralization, already during the mere instruction phase.

## 4 General Discussion

The present study investigated the neural and representational dynamics at play during S-R associative learning in different learning modes. We were first of all interested in extending previous research on instructed rapid learning to the investigation of the instruction period as opposed to the first implementation trials after instruction. Furthermore, we took into examination how this compares to learning by trial-and-error. Via the inclusion of a ‘mid-way condition’ that shared features with both instruction-based and trial-and-error learning, we were able to dissociate motor preparation from motor implementation and better understand the role that these play during learning.

In the following section, we initially explore the advantage of instruction-related behavior in learning compared to the other learning modes, focusing on the individual motor response representations observed during the instruction period in the absence of overt motor response implementation. Subsequently, we delve into the fMRI univariate analysis findings, which revealed a common frontoparietal decrease and DMN increase across conditions during learning, along with signal changes specific to instruction-based and trial-and-error learning trials. This is examined in the context of cognitive control demand. Lastly, we examine the findings from the ROI-based MVPA in prefrontal, parietal, and premotor ROIs, particularly discussing the consistency of brain representations across most ROIs in terms of early representations guiding later task performance.

### 4.1 The advantage of explicit instructions on learning: early proceduralization of instructed S-R rules

Behaviorally, we confirmed previous findings on the advantage during later task execution that explicit instructions offer when learning novel S-R rules as compared to rules that are explored via trial-and-error (and, in our design, via observation of correct and incorrect mappings). This, as previously assumed, points towards a detrimental effect of error processing, prominent in our trial-and-error and observation-based learning conditions, on learning successive correct S-R rules (Li et al., 2011; Monsell & Graham, 2021; cf. Ruge, Karcz, et al., 2018). Furthermore, it suggests that overt response implementation in trial-and-error learning, as opposed to merely observing correct and incorrect rules and feedback, mitigates the negative effect of error processing during learning on the subsequent implementation trials. To elaborate, both response accuracy as well as response speed in implementation trials in the trial-and-error condition showed values that fell between those observed in the instruction-based learning condition, where performance was at its peak, and the observation-based learning condition, where performance was at its lowest (Figure 4a and 4b). In our experimental design, the key distinction between learning via trial-and-error and mere observation was the requirement for overt response implementation in the former but not the latter. Therefore, we reasoned that the overall better performance in trial-and-error implementation trials, in contrast to previously observed implementation trials, may be attributed to learning via overt repeated attempts rather than the passive observation of correct and incorrect S-R links and feedback. The mechanism underlying this advantage of overtly implementing vs. merely observing correct and incorrect links, as hypothesized in the introduction, might involve a representational update in declarative as well as procedural WM at each incorrect feedback following an incorrectly implemented S-R link. This representational update could serve as a reset mechanism, potentially aiding the learning process. This hypothesis is supported by the significantly greater pattern similarity effect in trial-and-error compared to observed trials, as observed in the stage-specific pattern analysis of the left motor ROI (Figure 12a). Upon closer examination, we found that the pattern similarity effect in observed learning trials was not statistically significant as opposed to the other two learning conditions. This suggests that representations of individual motor programs are not represented in the left motor cortex during passive observation of correct and incorrect rules and feedback. Instead, our results suggest that for motor representations to be activated within the motor cortex when processing incorrect trials for learning, overt response implementation is a necessary condition. This contrasts with the instruction-based learning condition, where only correct S-R links were provided during learning. The stage-specific pattern analysis, in fact, revealed a significant pattern similarity effect in the left motor ROI already during instructed learning trials and in the absence of overt response implementation. This suggests that covert motor preparatory mechanisms are at play early during instructed learning trials and before response implementation, which in turn might explain the behavioral advantage of instruction-based learning over the other learning modes, especially in light of the significantly faster RTs in implementation trials (Figure 4b). Notably, motor representations observed during instructed learning trials were found in the same motor ROI that contained motor representations during implementation trials. This suggests that preparatory mechanisms during instruction-based learning occur in the same left motor regions where motor programs are represented during implementation trials (cf. Ariani et al., 2022). If during implementation trials the left motor cortex pattern similarity effect is likely to reflect overt motor implementation, on the other hand, during instructed learning trials it is plausible to assume that this mirrored covert motor implementation and motor imagery. The presence of motor representations during instructed learning trials in the same motor region that contained information during implementation trials is further complemented by the overlap that these representations showed in the cross-stage consistency analysis. These findings together suggest that under explicit instructions, procedural representations are formed as early as during learning and before implementation in the same motor regions that will later support response implementation, possibly guiding later performance during the implementation stage.

Last, it is important to highlight that a significantly stronger representational consistency between the learning and the implementation stage was observed when overt motor implementation was required for learning, as in the trial-and-error learning condition. This suggests that a stronger representational consistency between the learning and the implementation stage requires overt response execution, and that covert vs. overt response implementation leads to motor representations that are less consistent with those observed in the implementation stage.

### 4.2 Mean neural change during learning reflects decreasing cognitive control demand

The univariate analysis confirmed previous findings showing that key regions of the DMN increased in activity early during learning and across all learning modes. The DMN increase during learning was accompanied by a frontoparietal decrease along the inferior parietal sulcus and frontal midline, possibly mirroring lower cognitive control demand as learning proceeded. During implementation trials, the decrease was constrained to motor regions including postcentral gyrus/inferior parietal, vermis and cerebellum VI, left thalamus, and putamen, possibly supporting mechanisms related to visuo- and somatomotor adaptation (e.g. Floyer-Lea & Matthews, 2004; Makino et al., 2016 for a review).

#### 4.2.1 The role of the caudate head/ventral striatum in instruction-based learning trials

When inspecting linear signal change per each stimulus repetition that was specific to one learning mode as compared to the others, we found activity patterns specific to instruction and trial-and-error, but not to observed trials. One reason could logically be that our observation-based learning condition shared features with both instruction-based- and trial-and-error learning. Thus, it may be plausible to assume that observation-based learning in our paradigm was supported by neural mechanisms common to both instruction-based and trial-and-error learning.

Consistent with Ruge et al. (2019), where the ventral striatum was found to increase from the first implementation trial after instruction, we observed a similar signal increase in the instruction-based learning condition. However, in the present study, we further showed that this increase occurs early during learning, in the absence of overt response implementation. Additionally, we confirmed the specificity of this signal change to learning via explicit instructions in contrast with trial-and-error and observation-based learning.

The ventral striatum has been linked to reward processing, including positive feedback (Daniel & Pollmann, 2014 for an overview; e.g. Schott et al., 2007), and novel rule formation (Bédard & Sanes, 2009). In line with this, we observed relatively high activity in this region during trial-and-error learning trials, even at the initial repetition levels during learning. We hypothesized that this heightened activity in the ventral striatum during trial-and-error learning may be a result of immediate responsiveness to feedback-driven reward processing, especially when the correct response is overtly executed, as is the case in trial-and-error learning. This is in contrast to observed and instructed learning trials, where no overt execution of responses was required during the learning phase and, consequently, where we observed lower initial ventral striatal activity. During instructed learning trials, although no feedback is provided, we reasoned that covert response implementation, as indicated by the pattern similarity effect in the left motor ROI, likely occurs more frequently as the learner encounters the stimulus repeatedly. Thus, this gradual increase in covert response implementation might drive the incremental rise in ventral striatal activity from one repetition to the next that we observed in instructed trials (cf. Pasupathy & Miller, 2005).

Other regions that exhibited greater increases during instruction than during observation and trial-and-error learning included extended clusters in the right hemisphere, encompassing important hubs of the DMN such as the angular gyrus, precuneus, and ventromedial prefrontal cortex (VMPFC). We postulate that the overall simplicity of attending explicit instructions, coupled with the absence of ongoing interference caused by error trials in comparison to the other learning conditions, potentially led to a more rapid decrease in cognitive control demand. This reduction in demand likely manifested as a greater engagement of DMN-hub regions, which in our paradigm was more apparent in the right hemisphere under the instruction condition. This contrasts with the common bilateral DMN engagement observed across learning modes. Consistent with this rationale, when examining regions that exhibited a greater decrease in activity during trial-and-error learning compared to instructed and observed learning trials, results implicated regions associated with cognitive control demand, such as the DLPFC and anterior insula (e.g. Molnar-Szakacs & Uddin, 2022; Wu et al., 2019). Notably, these regions displayed relatively lower activity levels during instructed trials (Figure 7c), supporting the hypothesis that learning through explicit instructions places less cognitive control demand.

#### 4.2.2 Cingulo-opercular network and active rule exploration in trial-and-error learning

During trial-and-error learning trials, as opposed to the other conditions, we observed a greater activity decrease in the bilateral anterior insula (AI) and the bilateral anterior portion of the mid-cingulate cortex (aMCC, in the literature often referred to as dorsal or supracallosal anterior cingulate cortex; Figure 7b). These regions commonly co-activate as key components of the cingulo-opercular network (CON), known for its role in goal-directed cognitive control (Dosenbach et al., 2007; Dosenbach et al., 2008; Molnar-Szakacs & Uddin, 2022; Wood & Nee, 2023; Wu et al., 2019). The AI and the aMCC are suggested to support intentional action (Brass & Haggard, 2010; Hoffstaedter et al., 2014), evaluating action-outcomes to reinforce behavioral responses associated with more desirable results. One hypothesis is that the AI, with its rich connectivity to sensory areas (Dionisio et al., 2019; Gogolla, 2017), generates a “salience map” (Michel, 2017) of behaviorally relevant events (Brass & Haggard, 2010; Han et al., 2019; Hoffstaedter et al., 2014; Menon & Uddin, 2010; Touroutoglou et al., 2020) towards which attentional and WM resources (Hausman et al., 2021) can be directed for further processing and response (via the mACC, Han et al., 2019; Molnar-Szakacs & Uddin, 2022; Touroutoglou et al., 2020). This assessment influences the ability to transition between tasks and rules (Dosenbach et al., 2007; Dosenbach et al., 2008; Hausman et al., 2021; Menon & Uddin, 2010). In trial-and-error learning, this translates into evaluating stimulus-response-feedback and shifting between different responses for the same stimulus until the correct S-R mapping is implemented. This attentional shift ultimately contributes to forming and adjusting behavioral strategies, crucial for successful trial-and-error learning (Mohr et al., 2018). We propose that in trial-and-error learning, the CON supports the disengagement from previous error trials to facilitate the shift towards new S-R mappings, as well as the formation and implementation of adaptive and efficient learning strategies (Mohr et al., 2018). Importantly, according to our hypothesis, conditions that do not require to shift attentional resources from one mapping to another for the same stimulus, or necessitate response implementation and subsequent behavioral adjustment, should not elicit a comparable signal pattern in the CON-associated areas. In line with this, the instruction-based learning condition showed low and stable activity in both AI and aMCC (Figure 7c). The observation-based learning condition, on the other hand, plausibly required a representational shift akin to trial-and-error. However, the passive observation, as opposed to the active exploration of correct and incorrect rules during trial-and-error learning, plausibly mitigated the CON engagement, leading to lower initial activity in these regions (Figure 7c). Consistent with this reasoning, conditions that did not require motor implementation showed low activity in the aMCC, which supports top-down intentional movement with its connection to premotor, prefrontal, and parietal areas, as well as the cerebellum (Dosenbach et al., 2007; Hoffstaedter et al., 2014). Notably, these regions also decreased in activity during trial-and-error learning, possibly indicating that the activity in these areas decreased as a result of task automatization as learning progressed.

We considered two additional trial-and-error features that possibly drove a greater initial CON engagement as well as a subsequent signal decrease in trial-and-error learning. CON-related regions are sensitive to the level of uncertainty (or entropy, Fan, 2014; Wu et al., 2019), which plausibly was maximal at the beginning of trial-and-error learning, before any motor response was implemented, and gradually decreased via active rule exploration and the convergence towards predictable and accurate S-R mappings (Mohr et al., 2018). Importantly, this contrasts with the observation-based learning condition, which plausibly did not trigger the same level of uncertainty as trial-and-error learning because participants were not actively engaged in the motor response or the decision-making process. In other words, trial-and-error learning was the only condition in our study requiring context-dependent behavioral adjustment, which is supported by the CON as previously explained. Finally, AI and aMCC are sensitive to the rate of information under time constraints (Wu et al., 2019). We reasoned that the rate of information processing in trial-and-error learning plausibly was likely heightened as participants needed to rapidly cycle through the different previously incorrect implemented responses to identify (and implement within a certain time frame) the correct mapping. One hypothesis is therefore that the additional working-memory resources required for retrieving previous incorrect trials, coupled with the motor response requirement in trial-and-error learning, elicited a heightened CON activity and subsequent decrease in this condition.

### 4.3 Representational dynamics in prefrontal, parietal and premotor cortex

Our approach to the multivariate analysis was twofold. First, we aimed to identify brain regions where a pattern similarity effect could be detected across learning modes and stages. Once we established that individual rules were indeed represented in the brain, we employed an ROI-based approach. This enabled us to delve into more fine-grained questions about differences between regions, conditions, and stages. Importantly, the present study built upon Ruge et al. (2019) in extending the investigation of first implementation trials after instruction to the exploration of the initial learning phase. A key objective was to track the representational dynamics during S-R learning and implementation and assess the similarity between these representations as they transitioned from the learning to the implementation stage. Notably, this particular analysis was not feasible in the design of Ruge et al. (2019). To elaborate, finding rule information within the same brain region during both the learning and implementation stages, when pattern similarities are calculated separately within each stage, does not imply that these representations are similar. To address this, we employed a cross-stage consistency analysis, which allowed us to test for overlapping representations between learning and implementation trials in two distinct scenarios. The first scenario, exemplified by the trial-and-error condition, pertains to cases where response implementation was required early in the learning process. In contrast, the second scenario applies to learning conditions that did not require response implementation during the learning phase. In these instances, the transition to implementation trials was more profound and can be viewed as the shift from non-executed to executed trials. This latter scenario was particularly relevant to our instruction- and observation-based learning condition.

Rule information was identified in all predefined ROIs, encompassing the bilateral prefrontal and parietal cortices, as well as the bilateral premotor and left motor cortices. The presence of S-R rule information in frontoparietal regions is consistent with the role of these areas in exerting sustained attention and actively holding relevant information in WM (Chai et al., 2018; Cole et al., 2013; Dosenbach et al., 2007; Dosenbach et al., 2008; Ekman et al., 2016; Marvel et al., 2019). We observed stronger pattern similarity effects during learning as compared to implementation trials in prefrontal and parietal ROIs. This fits well with the typical frontoparietal mean activity decrease as learning progresses that we observed in the present study as well as in previous investigations (Bédard & Sanes, 2009; Hampshire et al., 2016, 2019; Mohr et al., 2016; Ruge & Wolfensteller, 2010, 2013).

During implementation trials, the pattern similarity effect in both the VLPFC and DLPFC did not reach statistical significance. This contrasts with Ruge et al. (2019), in which individual rules were found to be represented specifically in the left VLPFC during early implementation trials following instruction. However, our design contrasts with that of Ruge et al. (2019) in the way we tracked the whole learning stage. In fact, in Ruge et al. (2019) instructions were presented once and they were directly followed by implementation trials. Hence, it is plausible to assume that participants were still consolidating the one-time instructed rules at the first implementation trials after first-time instruction. This contrasts with our experimental design where each instructed S-R link was presented 4 times during the learning stage. Therefore, by the start of the implementation stage, rules were already learned, as apparent from the high behavioral accuracy in implementation trials. This might explain the low pattern similarity effect in prefrontal regions during implementation trials. In the parietal cortex, rules were more strongly represented in the superior part, which has been previously linked to information manipulation in WM (Crone et al., 2006; Koch et al., 2005; Koenigs et al., 2009; Wager & Smith, 2003). Notably, both the superior and inferior parts of the parietal cortex, as well as the VLPFC, exhibited overlapping rule representations across both the learning and implementation stages. This observation implies that these regions play a crucial role in constructing initial task representations (cf. Cole et al., 2011; González-García et al., 2017; Muhle-Karbe et al., 2017; Woolgar et al., 2016; Yang et al., 2019) during early learning trials, whether they involve overt execution as in trial-and-error or not as in instruction- and observation-based learning. Moreover, they seem to maintain this information consistently during rule execution, potentially aiding performance in subsequent rule implementation. This hypothesis was already proposed by Ruge et al. (2019). However, as explained before, representational consistency testing was not possible in the previous design. We, therefore, confirmed the early formation of abstract rule representation in VLPFC and DLPFC, as well as in the parietal cortex, and the maintenance of these very same representations throughout S-R implementation.

In the premotor ROIs, a significantly greater pattern similarity effect in learning vs. implementation trials was uniquely observed in the right vPMC/IFJ. This region is associated with abstract rule representation and rule updating (Brass & von Cramon, 2002, 2004; Derrfuss et al., 2004, 2005; Muhle-Karbe et al., 2017). Accordingly, the right IFJ has been found to contain decodable rule information during the delay period following instructions that had to be implemented later on as opposed to instructions that had to be simply memorized (Muhle-Karbe et al., 2017). This, together with the IFJ’s selective response to instruction cues before task implementation (Brass & von Cramon, 2002, 2004) suggests a role of this region in task preparation and might explain the greater similarity effect during learning vs. implementation trials. Individual rules were similarly represented across stages in the dPMC, a region associated with motor imagery (Pilgramm et al., 2016; Zabicki et al., 2019) as well as movement planning and execution (Hoshi & Tanji, 2007). Rule information in these regions likely mirrors preparatory mechanisms in the presence and absence of motor implementation during learning and overt motor execution during rule implementation. The premotor individual rule representations were similar from the learning to the implementation stages, suggesting that the premotor cortex is not only involved in preparatory mechanisms early during learning but maintains rule information during execution, possibly supporting rule implementation. Notably, the premotor cortex has been associated with information rehearsal in WM and it has been proposed to support the WM system beyond the traditional prefrontal and parietal regions (Marvel et al., 2019). This is consistent with the overlap of individual rule representations not only in frontoparietal but also in premotor cortices, supporting the idea that all these regions collectively contribute to maintaining rule information actively from learning to rule execution, possibly to guide task performance.

## 5 Conclusions

In the present study, we built upon prior investigations that focused on the initial implementation trials following first-time instruction. With a focus on the initial learning of S-R links via instruction and trial-and-error, we replicated and extended established results regarding the reorganization of brain activity in frontoparietal and DMN-related regions during learning (and not only implementation) trials. Crucially, the novelty of our study particularly lies in the exploration of individual rule identity representations during the critical transition from learning to implementation. Our study’s innovation lies in uncovering the early emergence of individual rule representations in prefrontal, parietal, premotor, and left motor cortices. Notably, these early representations align with those observed during subsequent implementation trials. This compelling observation suggests that abstract and procedural representations take shape during the early stages of learning, even in the absence of overt motor execution, and persist throughout motor implementation, possibly playing a crucial role in guiding successful task performance.

## Footnotes

1 Copyright Waiver. The above boilerplate text was automatically generated by fMRIPrep with the express intention that users should copy and paste this text into their manuscripts *unchanged*. It is released under the CC0 license.

## References

Abraham, A., Pedregosa, F., Eickenberg, M., Gervais, P., Mueller, A., Kossaifi, J., Gramfort, A., Thirion, B., & Varoquaux, G. (2014). Machine learning for neuroimaging with scikit-learn. Frontiers in Neuroinformatics, 8. 10.3389/fninf.2014.00014

Ariani, G., Pruszynski, J. A., & Diedrichsen, J. (2022). Motor planning brings human primary somatosensory cortex into action-specific preparatory states. eLife, 11, e69517. 10.7554/eLife.69517

Avants, B. B., Epstein, C. L., Grossman, M., & Gee, J. C. (2008). Symmetric diffeomorphic image registration with cross-correlation: Evaluating automated labeling of elderly and neurodegenerative brain. Medical Image Analysis, 12 (1), 26–41. 10.1016/j.media.2007.06.004

Baumann, A. W., Schäfer, T. A. J., & Ruge, H. (2023). Instructional load induces functional connectivity changes linked to task automaticity and mnemonic preference. NeuroImage, 277, 120262. 10.1016/j.neuroimage.2023.120262

Bédard, P., & Sanes, J. N. (2009). On a basal ganglia role in learning and rehearsing visual–motor associations. NeuroImage, 47 (4), 1701–1710. 10.1016/j.neuroimage.2009.03.050

Bode, S., & Haynes, J.-D. (2009). Decoding sequential stages of task preparation in the human brain. NeuroImage, 45 (2), 606–613. 10.1016/j.neuroimage.2008.11.031

Braem, S., Deltomme, B., & Liefooghe, B. (2019). The instruction-based congruency effect predicts task execution efficiency: Evidence from inter- and intra-individual differences. Memory & Cognition, 47 (8), 1582–1591. 10.3758/s13421-019-00951-3

Brass, M., & Haggard, P. (2010). The hidden side of intentional action: The role of the anterior insular cortex. Brain Structure and Function, 214 (5), 603–610. 10.1007/s00429-010-0269-6

Brass, M., Liefooghe, B., Braem, S., & De Houwer, J. (2017). Following new task instructions: Evidence for a dissociation between knowing and doing. Neuroscience and Biobehavioral Reviews, 81, 16–28. 10.1016/j.neubiorev.2017.02.012

Brass, M., & von Cramon, D. Y. (2002). The role of the frontal cortex in task preparation. Cerebral Cortex, 12 (9), 908–914. 10.1093/cercor/12.9.908

Brass, M., & von Cramon, D. Y. (2004). Selection for cognitive control: A functional magnetic resonance imaging study on the selection of task-relevant information. The Journal of Neuroscience: The Official Journal of the Society for Neuroscience, 24 (40), 8847–8852. 10.1523/JNEUROSCI.2513-04.2004

Brass, M., Wenke, D., Spengler, S., & Waszak, F. (2009). Neural Correlates of Overcoming Interference from Instructed and Implemented Stimulus–Response Associations. Journal of Neuroscience, 29 (6), 1766–1772. 10.1523/JNEUROSCI.5259-08.2009

Chai, W. J., Abd Hamid, A. I., & Abdullah, J. M. (2018). Working Memory From the Psychological and Neurosciences Perspectives: A Review. Frontiers in Psychology, 9. https://www.frontiersin.org/articles/10.3389/fpsyg.2018.00401

Cole, M. W., Bagic, A., Kass, R., & Schneider, W. (2010). Prefrontal dynamics underlying rapid instructed task learning reverse with practice. The Journal of Neuroscience: The Official Journal of the Society for Neuroscience, 30 (42), 14245–14254. 10.1523/JNEUROSCI.1662-10.2010

Cole, M. W., Etzel, J., Zacks, J., Schneider, W., & Braver, T. (2011). Rapid Transfer of Abstract Rules to Novel Contexts in Human Lateral Prefrontal Cortex. Frontiers in Human Neuroscience, 5. https://www.frontiersin.org/articles/10.3389/fnhum.2011.00142

Cole, M. W., Reynolds, J. R., Power, J. D., Repovs, G., Anticevic, A., & Braver, T. S. (2013). Multi-task connectivity reveals flexible hubs for adaptive task control. Nature Neuroscience, 16 (9, 9), 1348–1355. 10.1038/nn.3470

Collins, A. G. E., & Frank, M. J. (2013). Cognitive control over learning: Creating, clustering and generalizing task-set structure. Psychological Review, 120 (1), 190–229. 10.1037/a0030852

Cox, R. W., & Hyde, J. S. (1997). Software tools for analysis and visualization of fMRI data. NMR in Biomedicine, 10 (4-5), 171–178. 10.1002/(SICI)1099-1492(199706/08)10:4/5%3C171::AID-NBM453%3E3.0.CO;2-L

Crone, E. A., Wendelken, C., Donohue, S. E., & Bunge, S. A. (2006). Neural Evidence for Dissociable Components of Task-switching. Cerebral Cortex, 16 (4), 475–486. 10.1093/cercor/bhi127

Daniel, R., & Pollmann, S. (2014). A universal role of the ventral striatum in reward-based learning: Evidence from human studies. Neurobiology of Learning and Memory, 114, 90–100. 10.1016/j.nlm.2014.05.002

Derrfuss, J., Brass, M., Neumann, J., & von Cramon, D. Y. (2005). Involvement of the inferior frontal junction in cognitive control: Meta-analyses of switching and Stroop studies. Human Brain Mapping, 25 (1), 22–34. 10.1002/hbm.20127

Derrfuss, J., Brass, M., & von Cramon, D. Y. (2004). Cognitive control in the posterior frontolateral cortex: Evidence from common activations in task coordination, interference control, and working memory. NeuroImage, 23 (2), 604–612. 10.1016/j.neuroimage.2004.06.007

Dionisio, S., Mayoglou, L., Cho, S., Prime, D., Flanigan, P., Lega, B., Mosher, J., Leahy, R., Gonzalez-Martinez, J., & Nair, D. (2019). Connectivity of the human insula: A cortico-cortical evoked potential (CCEP) study. Cortex; a Journal Devoted to the Study of the Nervous System and Behavior, 120, 419–442. 10.1016/j.cortex.2019.05.019

Dosenbach, N. U. F., Fair, D. A., Cohen, A. L., Schlaggar, B. L., & Petersen, S. E. (2008). A dual-networks architecture of top-down control. Trends in Cognitive Sciences, 12 (3), 99–105. 10.1016/j.tics.2008.01.001

Dosenbach, N. U. F., Fair, D. A., Miezin, F. M., Cohen, A. L., Wenger, K. K., Dosenbach, R. A. T., Fox, M. D., Snyder, A. Z., Vincent, J. L., Raichle, M. E., Schlaggar, B. L., & Petersen, S. E. (2007). Distinct brain networks for adaptive and stable task control in humans. Proceedings of the National Academy of Sciences of the United States of America, 104 (26), 11073–11078. 10.1073/pnas.0704320104

Ekman, M., Fiebach, C. J., Melzer, C., Tittgemeyer, M., & Derrfuss, J. (2016). Different Roles of Direct and Indirect Frontoparietal Pathways for Individual Working Memory Capacity. Journal of Neuroscience, 36 (10), 2894–2903. 10.1523/JNEUROSCI.1376-14.2016

Entel, O., Tzelgov, J., & Bereby-Meyer, Y. (2014). Proportion congruency effects: Instructions may be enough. Frontiers in Psychology, 5, 1108. 10.3389/fpsyg.2014.01108

Esteban, O., Blair, R., Markiewicz, C. J., Berleant, S. L., Moodie, C., Ma, F., Isik, A. I., Erramuzpe, A., Kent, M., James D. and Goncalves DuPre, E., Sitek, K. R., Gomez, D. E. P., Lurie, D. J., Ye, Z., Poldrack, R. A., & Gorgolewski, K. J. (2018). fMRIPrep. Software. 10.5281/zenodo.852659

Esteban, O., Markiewicz, C., Blair, R. W., Moodie, C., Isik, A. I., Erramuzpe Aliaga, A., Kent, J., Goncalves, M., DuPre, E., Snyder, M., Oya, H., Ghosh, S., Wright, J., Durnez, J., Poldrack, R., & Gorgolewski, K. J. (2018). fMRIPrep: A robust preprocessing pipeline for functional MRI. Nature Methods. 10.1038/s41592-018-0235-4

Fan, J. (2014). An information theory account of cognitive control. Frontiers in Human Neuroscience, 8. https://www.frontiersin.org/articles/10.3389/fnhum.2014.00680

Floyer-Lea, A., & Matthews, P. M. (2004). Changing Brain Networks for Visuomotor Control With Increased Movement Automaticity. Journal of Neurophysiology, 92 (4), 2405–2412. 10.1152/jn.01092.2003

Fonov, V., Evans, A., McKinstry, R., Almli, C., & Collins, D. (2009). Unbiased nonlinear average age-appropriate brain templates from birth to adulthood. NeuroImage, 47*, Supplement* *1*, S102. 10.1016/S1053-8119(09)70884-5

Formica, S., González-García, C., & Brass, M. (2020). The effects of declaratively maintaining and proactively proceduralizing novel stimulus-response mappings. Cognition, 201, 104295. 10.1016/j.cognition.2020.104295

Friston, K. J., Penny, W. D., & Glaser, D. E. (2005). Conjunction revisited. NeuroImage, 25 (3), 661–667. 10.1016/j.neuroimage.2005.01.013

Gogolla, N. (2017). The insular cortex. Current Biology, 27 (12), R580–R586. 10.1016/j.cub.2017.05.010

González-García, C., Arco, J. E., Palenciano, A. F., Ramírez, J., & Ruz, M. (2017). Encoding, preparation and implementation of novel complex verbal instructions. NeuroImage, 148, 264–273. 10.1016/j.neuroimage.2017.01.037

González-García, C., Formica, S., Wisniewski, D., & Brass, M. (2021). Frontoparietal action-oriented codes support novel instruction implementation. NeuroImage, 226, 117608. 10.1016/j.neuroimage.2020.117608

Gorgolewski, K. J., Auer, T., Calhoun, V. D., Craddock, R. C., Das, S., Duff, E. P., Flandin, G., Ghosh, S. S., Glatard, T., Halchenko, Y. O., Handwerker, D. A., Hanke, M., Keator, D., Li, X., Michael, Z., Maumet, C., Nichols, B. N., Nichols, T. E., Pellman, J., . . . Poldrack, R. A. (2016). The brain imaging data structure, a format for organizing and describing outputs of neuroimaging experiments. Scientific Data, 3 (1, 1), 160044. 10.1038/sdata.2016.44

Gorgolewski, K. J., Burns, C. D., Madison, C., Clark, D., Halchenko, Y. O., Waskom, M. L., & Ghosh, S. (2011). Nipype: A flexible, lightweight and extensible neuroimaging data processing framework in python. Frontiers in Neuroinformatics, 5, 13. 10.3389/fninf.2011.00013

Gorgolewski, K. J., Esteban, O., Markiewicz, C. J., Ziegler, E., Ellis, D. G., Notter, M. P., Jarecka, D., Johnson, H., Burns, C., Manhães-Savio, A., Hamalainen, C., Yvernault, B., Salo, T., Jordan, K., Goncalves, M., Waskom, M., Clark, D., Wong, J., Loney, F., . . . Ghosh, S. (2018). Nipype. Software. 10.5281/zenodo.596855

Greve, D. N., & Fischl, B. (2009). Accurate and robust brain image alignment using boundary-based registration. NeuroImage, 48 (1), 63–72. 10.1016/j.neuroimage.2009.06.060

Hampshire, A., Daws, R. E., Neves, I. D., Soreq, E., Sandrone, S., & Violante, I. R. (2019). Probing cortical and sub-cortical contributions to instruction-based learning: Regional specialisation and global network dynamics. NeuroImage, 192, 88–100. 10.1016/j.neuroimage.2019.03.002

Hampshire, A., Hellyer, P. J., Parkin, B., Hiebert, N., MacDonald, P., Owen, A. M., Leech, R., & Rowe, J. (2016). Network mechanisms of intentional learning. NeuroImage, 127, 123–134. 10.1016/j.neuroimage.2015.11.060

Han, S. W., Eaton, H. P., & Marois, R. (2019). Functional Fractionation of the Cingulo-opercular Network: Alerting Insula and Updating Cingulate. Cerebral Cortex, 29 (6), 2624–2638. 10.1093/cercor/bhy130

Hartstra, E., Kühn, S., Verguts, T., & Brass, M. (2011). The implementation of verbal instructions: An fMRI study. Human Brain Mapping, 32 (11), 1811–1824. 10.1002/hbm.21152

Hausman, H. K., Hardcastle, C., Albizu, A., Kraft, J. N., Evangelista, N. D., Boutzoukas, E. M., Langer, K., O’Shea, A., Van Etten, E. J., Bharadwaj, P. K., Song, H., Smith, S. G., Porges, E., DeKosky, S. T., Hishaw, G. A., Wu, S., Marsiske, M., Cohen, R., Alexander, G. E., & Woods, A. J. (2021). Cingulo-opercular and frontoparietal control network connectivity and executive functioning in older adults. GeroScience, 44 (2), 847–866. 10.1007/s11357-021-00503-1

Hoffstaedter, F., Grefkes, C., Caspers, S., Roski, C., Palomero-Gallagher, N., Laird, A. R., Fox, P. T., & Eickhoff, S. B. (2014). The role of anterior midcingulate cortex in cognitive motor control: Evidence from functional connectivity analyses. Human Brain Mapping, 35 (6), 2741–2753. 10.1002/hbm.22363

Hoshi, E., & Tanji, J. (2007). Distinctions between dorsal and ventral premotor areas: Anatomical connectivity and functional properties. Current Opinion in Neurobiology, 17 (2), 234–242. 10.1016/j.conb.2007.02.003

Jenkinson, M., Bannister, P., Brady, M., & Smith, S. (2002). Improved optimization for the robust and accurate linear registration and motion correction of brain images. NeuroImage, 17 (2), 825–841. 10.1006/nimg.2002.1132

Jenkinson, M., & Smith, S. (2001). A global optimisation method for robust affine registration of brain images. Medical Image Analysis, 5 (2), 143–156. 10.1016/S1361-8415(01)00036-6

Koch, G., Oliveri, M., Torriero, S., Carlesimo, G. A., Turriziani, P., & Caltagirone, C. (2005). rTMS evidence of different delay and decision processes in a fronto-parietal neuronal network activated during spatial working memory. NeuroImage, 24 (1), 34–39. 10.1016/j.neuroimage.2004.09.042

Koenigs, M., Barbey, A. K., Postle, B. R., & Grafman, J. (2009). Superior Parietal Cortex Is Critical for the Manipulation of Information in Working Memory. Journal of Neuroscience, 29 (47), 14980–14986. 10.1523/JNEUROSCI.3706-09.2009

Lanczos, C. (1964). Evaluation of noisy data. Journal of the Society for Industrial and Applied Mathematics Series B Numerical Analysis, 1 (1), 76–85. 10.1137/0701007

Lee, S.-M., Henson, R. N., & Lin, C.-Y. (2020). Neural Correlates of Repetition Priming: A Coordinate-Based Meta-Analysis of fMRI Studies. Frontiers in Human Neuroscience, 14. https://www.frontiersin.org/articles/10.3389/fnhum.2020.565114

Lenth, R. V. (2023). Emmeans: Estimated marginal means, aka least-squares means. https://CRAN.R-project.org/package=emmeans

Li, J., Delgado, M. R., & Phelps, E. A. (2011). How instructed knowledge modulates the neural systems of reward learning. Proceedings of the National Academy of Sciences, 108 (1), 55–60. 10.1073/pnas.1014938108

Liefooghe, B., & De Houwer, J. (2018). Automatic effects of instructions do not require the intention to execute these instructions. Journal of Cognitive Psychology, 30, 108–121. 10.1080/20445911.2017.1365871

Liefooghe, B., Wenke, D., & De Houwer, J. (2012). Instruction-based task-rule congruency effects. *Journal of Experimental Psychology: Learning*, Memory, and Cognition, 38, 1325–1335. 10.1037/a0028148

Liu, X., Hairston, J., Schrier, M., & Fan, J. (2011). Common and distinct networks underlying reward valence and processing stages: A meta-analysis of functional neuroimaging studies. Neuroscience & Biobehavioral Reviews, 35 (5), 1219–1236. 10.1016/j.neubiorev.2010.12.012

Makino, H., Hwang, E. J., Hedrick, N. G., & Komiyama, T. (2016). Circuit Mechanisms of Sensorimotor Learning. Neuron, 92 (4), 705–721. 10.1016/j.neuron.2016.10.029

Marvel, C. L., Morgan, O. E., & Kronemer, S. I. (2019). How the Motor System Integrates with Working Memory. Neuroscience and Biobehavioral Reviews, 102, 184–194. 10.1016/j.neubiorev.2019.04.017

Meiran, N., Pereg, M., Kessler, Y., Cole, M. W., & Braver, T. S. (2015). Reflexive activation of newly instructed stimulus–response rules: Evidence from lateralized readiness potentials in no-go trials. Cognitive, Affective & Behavioral Neuroscience, 15 (2), 365–373. 10.3758/s13415-014-0321-8

Menon, V., & Uddin, L. Q. (2010). Saliency, switching, attention and control: A network model of insula function. Brain Structure and Function, 214 (5), 655–667. 10.1007/s00429-010-0262-0

Michel, M. (2017). A role for the anterior insular cortex in the global neuronal workspace model of consciousness. Consciousness and Cognition, 49, 333–346. 10.1016/j.concog.2017.02.004

Mohr, H., Wolfensteller, U., Betzel, R. F., Mišić, B., Sporns, O., Richiardi, J., & Ruge, H. (2016). Integration and segregation of large-scale brain networks during short-term task automatization. Nature Communications, 7 (1, 1), 13217. 10.1038/ncomms13217

Mohr, H., Zwosta, K., Markovic, D., Bitzer, S., Wolfensteller, U., & Ruge, H. (2018). Deterministic response strategies in a trial-and-error learning task. PLoS Computational Biology, 14 (11), e1006621. 10.1371/journal.pcbi.1006621

Molnar-Szakacs, I., & Uddin, L. Q. (2022). Anterior insula as a gatekeeper of executive control. Neuroscience & Biobehavioral Reviews, 139, 104736. 10.1016/j.neubiorev.2022.104736

Monsell, S., & Graham, B. (2021). Role of verbal working memory in rapid procedural acquisition of a choice response task. Cognition, 214, 104731. 10.1016/j.cognition.2021.104731

Muhle-Karbe, P. S., Duncan, J., De Baene, W., Mitchell, D. J., & Brass, M. (2017). Neural Coding for Instruction-Based Task Sets in Human Frontoparietal and Visual Cortex. Cerebral Cortex, 27 (3), 1891–1905. 10.1093/cercor/bhw032

Mumford, J. A., Davis, T., & Poldrack, R. A. (2014). The impact of study design on pattern estimation for single-trial multivariate pattern analysis. NeuroImage, 103, 130–138. 10.1016/j.neuroimage.2014.09.026

Mumford, J. A., Turner, B. O., Ashby, F. G., & Poldrack, R. A. (2012). Deconvolving BOLD activation in event-related designs for multivoxel pattern classification analyses. Neuroimage, 59 (3), 2636–2643. 10.1016/j.neuroimage.2011.08.076

Nichols, T., Brett, M., Andersson, J., Wager, T., & Poline, J.-B. (2005). Valid conjunction inference with the minimum statistic. NeuroImage, 25 (3), 653–660. 10.1016/j.neuroimage.2004.12.005

Oberauer, K. (2009). Chapter 2 Design for a Working Memory. In Psychology of Learning and Motivation (Vol. 51, pp. 45–100). Elsevier. 10.1016/S0079-7421(09)51002-X

Oberauer, K. (2010). Declarative and procedural working memory: Common principles, common capacity limits? Psychologica Belgica, 50 (3-4), 277–308. 10.5334/pb-50-3-4-277

Oosterhof, N. N., Connolly, A. C., & Haxby, J. V. (2016). CoSMoMVPA: Multi-Modal Multivariate Pattern Analysis of Neuroimaging Data in Matlab/GNU Octave. Frontiers in Neuroinformatics, 10, 27. 10.3389/fninf.2016.00027

Palenciano, A. F., González-García, C., de Houwer, J., Brass, M., & Liefooghe, B. (2021). Exploring the Link between Novel Task Proceduralization and Motor Simulation. Journal of Cognition, 4 (1), 57. 10.5334/joc.190

Pasupathy, A., & Miller, E. K. (2005). Different time courses of learning-related activity in the prefrontal cortex and striatum. Nature, 433 (7028), 873–876. 10.1038/nature03287

Pilgramm, S., de Haas, B., Helm, F., Zentgraf, K., Stark, R., Munzert, J., & Krüger, B. (2016). Motor imagery of hand actions: Decoding the content of motor imagery from brain activity in frontal and parietal motor areas. Human Brain Mapping, 37 (1), 81–93. 10.1002/hbm.23015

Raichle, M. E. (2015). The brain’s default mode network. Annual Review of Neuroscience, 38, 433–447. 10.1146/annurev-neuro-071013-014030

Ruge, H., Karcz, T., Mark, T., Martin, V., Zwosta, K., & Wolfensteller, U. (2018). On the efficiency of instruction-based rule encoding. Acta Psychologica, 184, 4–19. 10.1016/j.actpsy.2017.04.005

Ruge, H., Legler, E., Schäfer, T. A. J., Zwosta, K., Wolfensteller, U., & Mohr, H. (2018). Unbiased Analysis of Item-Specific Multi-Voxel Activation Patterns Across Learning. Frontiers in Neuroscience, 12. 10.3389/fnins.2018.00723

Ruge, H., Schäfer, T. A., Zwosta, K., Mohr, H., & Wolfensteller, U. (2019). Neural representation of newly instructed rule identities during early implementation trials. eLife, 8, e48293. 10.7554/eLife.48293

Ruge, H., & Wolfensteller, U. (2010). Rapid formation of pragmatic rule representations in the human brain during instruction-based learning. *Cerebral Cortex (New York*, N.Y.: 1991), *20* (7), 1656–1667. 10.1093/cercor/bhp228

Ruge, H., & Wolfensteller, U. (2013). Functional integration processes underlying the instruction-based learning of novel goal-directed behaviors. NeuroImage, 68, 162–172. 10.1016/j.neuroimage.2012.12.003

Ruge, H., & Wolfensteller, U. (2015). Distinct fronto-striatal couplings reveal the double-faced nature of response–outcome relations in instruction-based learning. *Cognitive, Affective*, & Behavioral Neuroscience, 15 (2), 349–364. 10.3758/s13415-014-0325-4

Schott, B. H., Niehaus, L., Wittmann, B. C., Schütze, H., Seidenbecher, C. I., Heinze, H.-J., & Düzel, E. (2007). Ageing and early-stage Parkinson’s disease affect separable neural mechanisms of mesolimbic reward processing. Brain, 130 (9), 2412–2424. 10.1093/brain/awm147

Singmann, H., Bolker, B., Westfall, J., Aust, F., & Ben-Shachar, M. S. (2023). Afex: Analysis of factorial experiments. https://CRAN.R-project.org/package=afex

Spunt, B. (2016). Spunt/bspmview: BSPMVIEW v.20161108. Zenodo. 10.5281/zenodo.168074

Touroutoglou, A., Andreano, J., Dickerson, B. C., & Barrett, L. F. (2020). The Tenacious Brain: How the Anterior Mid-Cingulate Contributes to Achieving Goals. Cortex; a Journal Devoted to the Study of the Nervous System and Behavior, 123, 12–29. 10.1016/j.cortex.2019.09.011

Tustison, N. J., Avants, B. B., Cook, P. A., Zheng, Y., Egan, A., Yushkevich, P. A., & Gee, J. C. (2010). N4ITK: Improved N3 bias correction. IEEE Transactions on Medical Imaging, 29 (6), 1310–1320. 10.1109/TMI.2010.2046908

Tzourio-Mazoyer, N., Landeau, B., Papathanassiou, D., Crivello, F., Etard, O., Delcroix, N., Mazoyer, B., & Joliot, M. (2002). Automated anatomical labeling of activations in SPM using a macroscopic anatomical parcellation of the MNI MRI single-subject brain. NeuroImage, 15 (1), 273–289. 10.1006/nimg.2001.0978

Wager, T. D., & Smith, E. E. (2003). Neuroimaging studies of working memory: *Cognitive, Affective*, & Behavioral Neuroscience, 3 (4), 255–274. 10.3758/CABN.3.4.255

Wenke, D., De Houwer, J., De Winne, J., & Liefooghe, B. (2015). Learning through instructions vs. Learning through practice: Flanker congruency effects from instructed and applied S-R mappings. Psychological Research, 79 (6), 899–912. 10.1007/s00426-014-0621-1

Wolfensteller, U., & Ruge, H. (2012). Frontostriatal mechanisms in instruction-based learning as a hallmark of flexible goal-directed behavior. Frontiers in Psychology, 3. 10.3389/fpsyg.2012.00192

Wood, J. L., & Nee, D. E. (2023). Cingulo-Opercular Subnetworks Motivate Frontoparietal Subnetworks during Distinct Cognitive Control Demands. Journal of Neuroscience, 43 (7), 1225–1237. 10.1523/JNEUROSCI.1314-22.2022

Woolgar, A., Jackson, J., & Duncan, J. (2016). Coding of Visual, Auditory, Rule, and Response Information in the Brain: 10 Years of Multivoxel Pattern Analysis. Journal of Cognitive Neuroscience, 28 (10), 1433–1454. 10.1162/jocn_a_00981

Wu, T., Wang, X., Wu, Q., Spagna, A., Yang, J., Yuan, C., Wu, Y., Gao, Z., Hof, P. R., & Fan, J. (2019). Anterior insular cortex is a bottleneck of cognitive control. NeuroImage, 195, 490–504. 10.1016/j.neuroimage.2019.02.042

Yang, G. R., Joglekar, M. R., Song, H. F., Newsome, W. T., & Wang, X.-J. (2019). Task representations in neural networks trained to perform many cognitive tasks. Nature Neuroscience, 22 (2, 2), 297–306. 10.1038/s41593-018-0310-2

Zabicki, A., de Haas, B., Zentgraf, K., Stark, R., Munzert, J., & Krüger, B. (2019). Subjective vividness of motor imagery has a neural signature in human premotor and parietal cortex. NeuroImage, 197, 273–283. 10.1016/j.neuroimage.2019.04.073

Zhang, Y., Brady, M., & Smith, S. (2001). Segmentation of brain MR images through a hidden markov random field model and the expectation-maximization algorithm. IEEE Transactions on Medical Imaging, 20 (1), 45–57. 10.1109/42.906424

## Software

Psychology Software Tools, Inc. [E-Prime 3.0]. (2016). Retrieved from https://support.pstnet.com/.

The Math Works, Natick, MA, United States. Matlab 9.9.0.1467703 (R2020b) (August 26, 2020). available online from http://www.mathworks.com

R Core Team (2021). R: A language and environment for statistical computing. R Foundation for Statistical Computing, Vienna, Austria. URL https://www.R-project.org/.

