## supplementary material for "Initial Learning in the Brain: From Rules to Action"

### Preregistration

The study was initially preregistered on AsPredicted (https://aspredicted.org/9q9m6.pdf). Along the study, we made some changes that are described below.

**Inclusion criteria.** We initially planned to exclude subjects whose accuracy fell below 75% during implementation trials and 50% during learning trials for at least one stimulus in the trial-and-error condition. The rationale behind this was to optimize trial-and-error blocks before yoking them to the observation blocks between participants. However, this would have required analyzing subjects' performance during data acquisition to address potential dropouts and constantly have an available data set for the trial-and-error-to-observation block yoking procedure. Upon evaluating the feasibility of analyzing subjects' behavior during data acquisition, we realized that it posed challenges in terms of scanner booking times and participant scheduling. Consequently, we adopted a less stringent inclusion criterion, which was set at an 80% overall accuracy rate across conditions and blocks. The overall accuracy was calculated during the experiment by E-Prime itself and did not therefore require analysis behavior during data acquisition. Nonetheless, we conducted the pre-registered accuracy analysis at the end of the data acquisition phase. We identified seven participants who did not meet the pre-registered inclusion criteria for one to three stimuli in 1 out of 8 trial-and-error learning blocks (mean stimuli per block = 2.14, SD = 0.37). Subsequently, we examined the performance of these participants who were yoked in their observation-based learning blocks to those who did not perform well in the trial-and-error blocks. Surprisingly, all 'yoked' participants were able to successfully learn the S-R links.

In more detail, while participants from whom the response patterns were taken scored less than 50% and 75% hits in trial-and-error learning and implementation trials, respectively, the ‘observation-yoked’ participants met the 50-75 inclusion criterion. An example is provided in Table S1 for the observation condition (i.e., yoked participant) and S2 for the trial-and-error condition (i.e., source participant). Only one participant did not meet the 50-75 inclusion criterion in one observation block for 50% of the presented stimuli. However, in line with our approach of not excluding any other subjects or blocks analyzed within the scope of the 50-75 criterion, we decided to include this subject and block as well.

**ROI-based MVPA contrasts.** The planned stage-specific contrasts for repetition levels 1-2 vs. 3-4 and 5-6 vs. 7-8, as well as cross-stage consistency comparison of repetitions 3-4 vs. 5-6 were not implemented as pre-registered. Instead, the MVPA procedure was optimized to compute pairwise comparisons instead of mean values between repetitions of interest. This allowed for a more precise and reliable trial-by-trial analysis. For a more detailed description, refer to the MVPA analysis section.

**Connectivity analysis.** Given the length and complexity of the present paper, we have chosen to include the connectivity analysis in a separate paper.

Table S1

*Example of single-subject performance in the observation-based learning condition when the source subject of the trial-and-error condition performed poorly in the learning stage.*

| Condition | Sub | Run | Block | TE Source Sub | Trial | Stimulus | Accuracy |
|---|---|---|---|---|---|---|---|
| OBS | 13 | 4 | 7 | 8 | 3 | 1 | 0 |
| OBS | 13 | 4 | 7 | 8 | 8 | 1 | 0 |
| OBS | 13 | 4 | 7 | 8 | 10 | 1 | 0 |
| OBS | 13 | 4 | 7 | 8 | 16 | 1 | 1 |
| OBS | 13 | 4 | 7 | 8 | 19 | 1 | 1 |
| OBS | 13 | 4 | 7 | 8 | 21 | 1 | 1 |
| OBS | 13 | 4 | 7 | 8 | 27 | 1 | 1 |
| OBS | 13 | 4 | 7 | 8 | 31 | 1 | 1 |
| OBS | 13 | 4 | 7 | 8 | 1 | 4 | 0 |
| OBS | 13 | 4 | 7 | 8 | 6 | 4 | 0 |
| OBS | 13 | 4 | 7 | 8 | 11 | 4 | 0 |
| OBS | 13 | 4 | 7 | 8 | 15 | 4 | 1 |
| OBS | 13 | 4 | 7 | 8 | 18 | 4 | 1 |
| OBS | 13 | 4 | 7 | 8 | 24 | 4 | 1 |
| OBS | 13 | 4 | 7 | 8 | 25 | 4 | 1 |
| OBS | 13 | 4 | 7 | 8 | 29 | 4 | 1 |

OBS = Observation-based learning condition.

Sub = Subject

TE Source Sub = the subject from whom the trial-and-error learning response patterns were collected as part of the yoking procedure.

Table S2

*Source subject's performance in the trial-and-error condition.*

| Condition | Sub | Run | Block | Trial | Stimulus | Accuracy |
|---|---|---|---|---|---|---|
| TE | 8 | 4 | 8 | 3 | 1 | 0 |
| TE | 8 | 4 | 8 | 8 | 1 | 0 |
| TE | 8 | 4 | 8 | 10 | 1 | 0 |
| TE | 8 | 4 | 8 | 16 | 1 | 1 |
| TE | 8 | 4 | 8 | 20 | 1 | 0 |
| TE | 8 | 4 | 8 | 23 | 1 | 0 |
| TE | 8 | 4 | 8 | 27 | 1 | 0 |
| TE | 8 | 4 | 8 | 30 | 1 | 0 |
| TE | 8 | 4 | 8 | 1 | 4 | 0 |
| TE | 8 | 4 | 8 | 6 | 4 | 0 |
| TE | 8 | 4 | 8 | 11 | 4 | 0 |
| TE | 8 | 4 | 8 | 15 | 4 | 1 |
| TE | 8 | 4 | 8 | 19 | 4 | 0 |
| TE | 8 | 4 | 8 | 24 | 4 | 0 |
| TE | 8 | 4 | 8 | 26 | 4 | 0 |
| TE | 8 | 4 | 8 | 31 | 4 | 0 |

TE = Trial-and-Error learning condition.

Sub = Subject
